# Distinct nanoscale calcium channel and synaptic vesicle topographies contribute to the diversity of synaptic function

**DOI:** 10.1101/421370

**Authors:** Nelson Rebola, Maria Reva, Tekla Kirizs, Miklos Szoboszlay, Gael Moneron, Zoltan Nusser, David A. DiGregorio

## Abstract

The nanoscale topographical arrangement of voltage-gated calcium channels (VGCC) and synaptic vesicles (SVs) determines synaptic strength and plasticity, but whether distinct spatial distributions underpin diversity of synaptic function is unknown. We performed single bouton Ca^2+^ imaging, Ca^2+^ chelator competition, immunogold electron microscopic (EM) localization of VGCCs and the active zone (AZ) protein Munc13-1, at two cerebellar synapses. Unexpectedly, we found that weak synapses exhibited 3-fold more VGCCs than strong synapses, while the coupling distance was 5-fold longer. Reaction-diffusion modelling could explain both functional and structural data with two strikingly different nanotopographical motifs: strong synapses are composed of SVs that are tightly coupled (∼10 nm) to VGCC clusters, whereas at weak synapses VGCCs were excluded from the vicinity (∼50 nm) of docked vesicles. The distinct VGCC-SV topographical motifs also confer differential sensitivity to neuromodulation. Thus VGCC-SV arrangements are not canonical across CNS synapses and their diversity could underlie functional heterogeneity.

## Introduction

Chemical synapses are thought to be fundamental computational units of neuronal networks, routing information throughout circuits, and are cellular substrates for learning and memory. Synapses transform electrical impulses (action potentials, APs) of one cell into electrical signals (postsynaptic potentials, PSPs) of another through the opening of VGCCs, which increase the intracellular [Ca^2+^] and trigger synaptic vesicle exocytosis of chemical neurotransmitters. This intricate cascade of signal transduction requires the organization of protein complexes (e.g. ion channels, membrane fusion proteins and receptors) and SVs at scale of tens of nanometers (Sudhof, 2012) to set synaptic strength (Biederer et al., 2017) and convey information with sub-millisecond temporal precision (Eggermann et al., 2012). But whether the precise arrangement of SVs and VGCCs is uniform across central synapses or its variability contributes to the diversity of synaptic function throughout the brain is unknown.

Synapses exhibit functional heterogeneity in strength and short-term plasticity (STP; Atwood and Karunanithi, 2002; Dittman et al., 2000; Grande and Wang, 2011; Millar et al., 2002) which are critical for circuit computations (Abbott and Regehr, 2004; Chabrol et al., 2015; Chadderton et al., 2014; Diaz-Quesada et al., 2014). The function of synapses can be quantified using analysis methods based on the quantal theory of transmission (Katz, 1969), where synaptic strength elicited by a single presynaptic AP can be estimated by the product of three parameters: number of release sites (N), the probability that a single vesicle releases neurotransmitter (P_v_), and the quantal size (q). The quantal size is largely influenced by the number of neurotransmitter-gated ion channels in the postsynaptic density. Our understanding of the physical basis of N within a single AZ is only recently starting to emerge with the observation of distinct calcium channel clusters driving SV fusion (Holderith et al., 2012; Miki et al., 2017; Nakamura et al., 2015) and the observation that release occurs from multiple locations, each marked by clusters of Munc13-1 (Sakamoto et al., 2018).

P_v_ not only influences synaptic strength, but also reliability and STP (Abbott and Regehr, 2004) and is known to be heterogeneous across CNS synapses (Chabrol et al., 2015; Dittman et al., 2000; Holderith et al., 2012; Koester and Johnston, 2005). While there is a plethora of molecules implicated in AP-triggered vesicle fusion (Korber and Kuner, 2016; Sudhof, 2012), very few are known to account for the diversity of P_v_. Variation in VGCC density and number within presynaptic clusters have been shown to drive heterogeneity of P_v_ (Koester and Johnston, 2005; Nakamura et al., 2015). Heterogeneity of P_v_ could also arise from variations in the distance between VGCCs and SVs. For example high P_v_ synapses are thought to exhibit short coupling distances (Bucurenciu et al., 2010; Rozov et al., 2001), while low P_v_ synapses exhibit longer distances (Rozov et al., 2001; Vyleta and Jonas, 2014). These distances are typically estimated from Ca^2+^ chelator competition experiments along with numerical simulations, but alone cannot predict the 2-dimensional arrangement of VGCCs and SVs (Nakamura et al., 2018).

To date specific VGCCs-SV topographies have been proposed at Drosophila (VGCC clusters coupled to vesicles around its perimeter) and mammalian NMJ (rows of VGCC and SVs), and at hair cell synapses (rows of calcium channels coupled to SV around its perimeter; Dittrich et al., 2013; Luo et al., 2015; Neef et al., 2018; Pangrsic et al., 2015). However, the nanoscale organization of SVs and VGCCs underlying the functional diversity of CNS synapses is less well understood. At the giant synapse of calyx of Held, ultrastructural reconstructions and numerical simulations argued for multiple docked vesicles per AZ and a random distribution of VGCCs (Keller et al., 2015). In a separate study at the same synapse SDS-digested freeze-fracture replica-immunolabeling (SDS-FRL) revealed that VGCCs appear in clusters much smaller than the AZ, which in combination with other biophysical experiments and mathematical modelling, argued that vesicles fused at sites within less than 20 nm from the perimeter of VGCCs clusters in mature terminals (Nakamura et al., 2015). Whether this model represents a canonical topographical motif, particularly within small bouton-type central synapses, is presently unknown. This is partly due to the inability to estimate the location of VGCCs and SVs synaptic vesicles at the same synapse.

Detailed studies of neurotransmitter release from typical boutons are challenging due to their small size. Here we take advantage of optical approaches to estimate the biophysical parameters of Ca^2+^ entry, a critical parameter influencing estimates of coupling distance (Nakamura et al., 2018), at two different small bouton-type synapses within acute cerebellar brain slices: one strong (formed by cerebellar stellate cells; SCs) and one weak (formed by cerebellar granule cells; GCs). In order to estimate the 2D arrangement of VGCCs and SV underlying synaptic function at both synapses, Ca^2+^ chelator-based coupling distance estimates were complimented with SDS-FRL measurements of VGCC and Munc13-1 (SV fusion sites) distributions and Monte Carlo simulations of buffered Ca^2+^ diffusion and binding to a release sensor. We provide the first evidence that the nanotopography of VGCCs and SVs is consistent with a clustered model in SCs, and in contrast, VGCCs in GCs are distributed uniformly throughout the AZ except in a region surrounding docked vesicles (exclusion zone, EZ).

## Results

### Amplitude of presynaptic Ca^2+^ transients do not correlate with P_v_

In cortical glutamatergic synapses, the amplitude of presynaptic Ca^2+^ transients is positively correlated with release probability (Holderith et al., 2012; Koester and Johnston, 2005). We therefore measured the amplitude of single AP-evoked Ca^2+^ transients (sAPCaTs) in boutons of weak GC to Purkinje cell (PC) synapses (P_v_ ∼ 0.2; Baur et al., 2015; Valera et al., 2012) and in boutons of strong SC-SC synapses (P_v_ 0.3-0.8; Arai and Jonas, 2014; Pulido et al., 2015). GC and SC were whole-cell patch-clamped with a pipette containing Alexa-594 (10 μM) and the Ca^2+^ indicator Fluo-5F (100 μM) in the internal solution (Figure 1A, C). We elicited sAPCaTs at specific times after establishing the whole-cell configuration: 15 minutes for GCs and 30 minutes for SCs, corresponding to ∼70% and ∼80% dialysis, respectively (Figure S1A-F). Unexpectedly, sAPCaTs recorded from low P_v_ GC boutons were four times larger than those recorded from the moderate P_V_ SC boutons (Figure 1E; GC: ΔF/F: 5.82 ± 0.46, n = 23 boutons, 4 cells; SC: ΔF/F: 1.37 ± 0.07, n = 70 boutons, 11 cells). Similar results were obtained using the low affinity Ca^2+^ indicator OGB-5N (Figure 1F). The larger average sAPCaT in SC boutons contrasts previous reports that sAPCaT amplitudes predict synaptic strength (Eltes et al., 2017; Holderith et al., 2012; Koester and Johnston, 2005).

**Figure 1:**
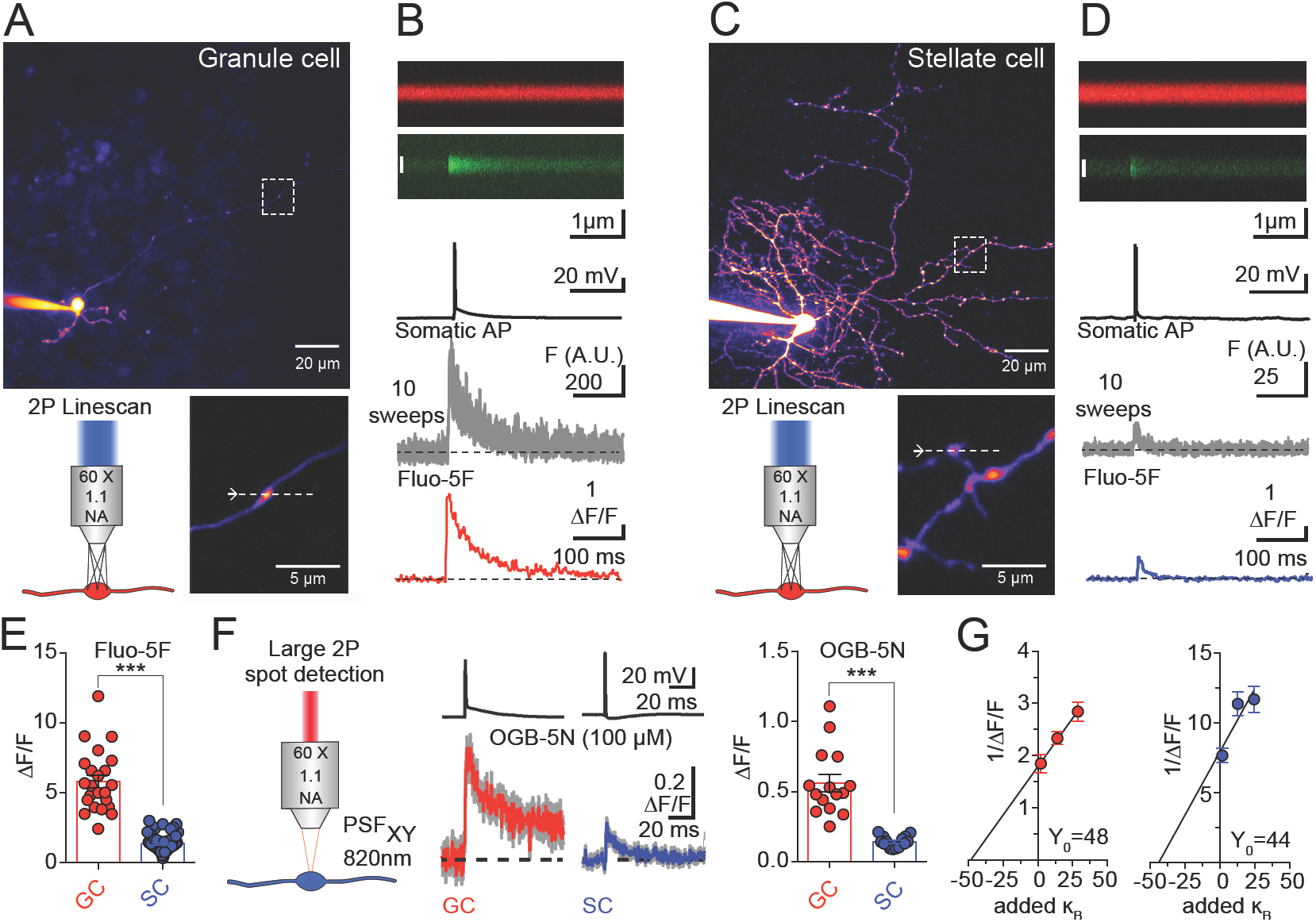
Single action potential-evoked Ca^2+^ transients (sAPCaTs) do not correlate with synaptic strength. **(A)** *Top*: 2PLSM image (maximum-intensity projection, MIP) of granule cell (GC) loaded with Alexa-594 (10 μM). *Bottom*: sAPCaTs were recorded using 2P line scan (dashed line) of single boutons (dashed box in upper image). **(B)** Averaged line scan image (from 10 images) of Alexa-594 (red) and Fluo-5F (green: 100 μM) fluorescence (top) recorded from bouton in A following AP initiation by current injection (500 pA, 0.5 ms). Time series traces are mean fluorescence of individual images over 0.7 μm (white line), and the red trace is the average of all 10 traces after conversion to ΔF/F. **(C-D)** Same as in **A** and **B**, but for SCs. **(E)** peak amplitude of sAPCaTs in GCs (n = 23 boutons) and SCs (n = 70). Box indicates mean ± SEM. (*** p<0.001, Mann-Whitney test). **(F)** *Left*: A large illumination spot was created by underfilling the objective pupil with a collimated laser beam (PSF_XY_: 0.82 ± 0.01 μm; PSF_z_: 10.2 ± 0.27 μm, n = 12). *Middle*: single AP recordings (top) and average (bottom; from 8 line scan sweeps) sAPCaTs recorded using large spot detection and a low affinity indicator OGB-5N (100 μM concentration). *Right:* sAPCaTs peak amplitudes for OGB-5N in GC (n = 16) and SC (n = 15) boutons. (*** p<0.001, Mann-Whitney test) **(G)** Inverse sAPCaTs amplitude plotted against Ca^2+^-binding ratio (κB) of the added indicator OGB-5N. The lines represent linear fits. Extrapolation to the abscissa gives an estimate of the Ca^2+^-binding ratio of endogenous fixed buffers.

We did not detect differences in the endogenous buffer capacity (GC: κ_E_= 47 ± 14; SCs: κ_E_= 44 ± 18; Figure 1G), and after correction for a 30% volume difference observed between SC and GC boutons (Figure S1G-I), a 3-fold difference in sAPCaT peak amplitude remained. We therefore conclude that larger sAPCaT amplitude in GCs arises from an increase in total Ca entry rather than from a decreased endogenous Ca buffering compared to SCs.

### GC boutons possess larger numbers of VGCCs

In order to determine the source of the difference in Ca^2+^ entry, fluctuation analysis (Sabatini and Svoboda, 2000) of sAPCaTs was used to estimate the VGCC number and open probability (P_open_). 60-100 consecutive sAPCaTs were extracellularly evoked following transient Ca^2+^ indicator loading (Figure 2A-B). Despite the large sAPCaTs in GC boutons, Fluo-5F linearly reported [Ca^2+^] changes (Figure S2A-B). To improve the signal-to-noise ratio of sAPCaT recordings from SC, we used the high affinity Ca^2+^ indicator Fluo-4, which also linearly reported [Ca^2+^] changes (Figure S2C). The coefficient of variation (CV) of the shot-noise subtracted trial-to-trial peak fluorescence was smaller in GC than in SC (Figure 2D-E).

**Figure 2:**
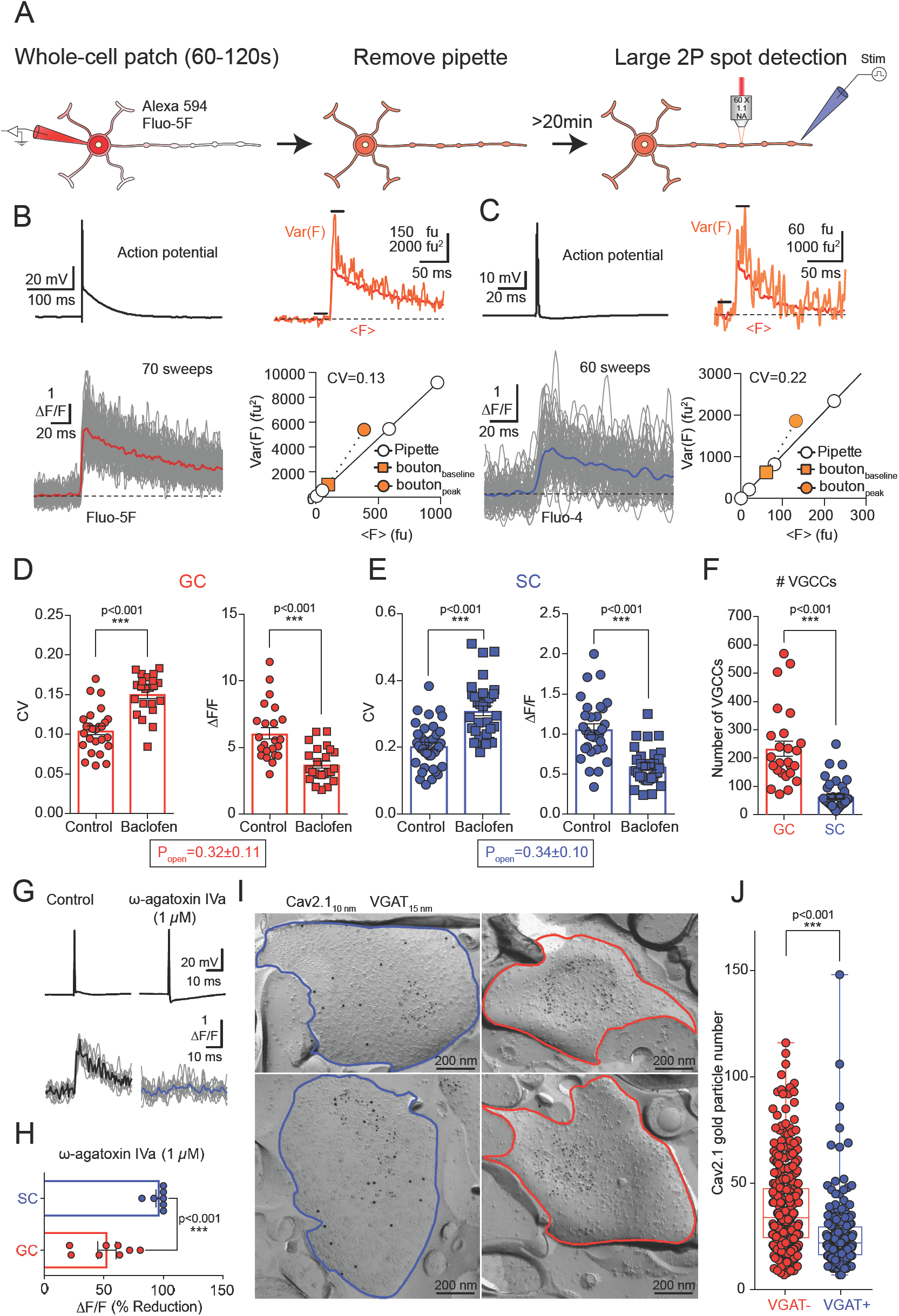
Estimation of total VGCCs number and open probability. **(A)** Schematic illustration showing brief indicator loading and extracellular stimulation to elicit sAPCaTs. **(B)** *Left*: single somatic AP (*top*) and corresponding Fluo-5F sAPCaTs in GC boutons (averaged trace: red; *bottom*). *Right*: Variance from the 70 recorded sAPCaTs compared to variance predicted by dark and shot noise alone (red trace). Mean variance plotted against mean (bottom right) for 20 ms baseline (orange square) and 20 ms following the peak of the fluorescence transient. Open circles, variance vs mean relationship obtained by imaging the patch pipette at different illumination intensities to estimate variance expected from photon shot noise. fu, fluorescence unit. **(C)** Same as in **B**, but with the Ca^2+^ indicator Fluo-4 in SCs **(D)** Summary plot of CV and peak amplitude of [Ca^2+^] transients obtained in control conditions and in the presence of 10 μM Baclofen for GC boutons. (*** p<0.001, Mann-Whitney test) **(E)** Same as in **D**, but for SCs. (*** p<0.001, Mann-Whitney test) **(F)** Summary plot for calculated number of VGCCs present in GC (n = 26) and SC (n = 39) boutons. (*** p<0.001, Mann-Whitney test) **(G)** Somatic AP (top) and recorded [Ca^2+^] transient (Fluo-5F, 100 μM) in corresponding SC bouton (bottom), in control conditions (left) and after the application of ω-agatoxin-IVA (1μM, right). **(H)** Summary plot of the effect of ω-agatoxin-IVA on the amplitude of sAPCaTs in SC and GC boutons. (*** p<0.001, Mann-Whitney test) **(I)** High-resolution SDS-FRL localization of Cav2.1 subunits (10 nm gold) in putative SC (VGAT-positive, 15 nm gold, left) and GC (VGAT-negative, right) boutons. Red and blue lines delineate bouton surface. **(J)** Summary plot of the number of immunogold particles per VGAT-positive (n = 184) and VGAT-negative (n = 426) boutons. (*** p<0.001, Mann-Whitney test)

The number and P_open_ of VGCCs was calculated from the mean and variance of the peak amplitude of sAPCaTs in the presence and absence of a neuromodulator (Sabatini and Svoboda, 2000). GABA_B_ receptor agonist baclofen reduced sAPCaT peak amplitude and simultaneously increased the CV both in SC and GC boutons (Figure 2D, E), consistent with a change in P_open_ of VGCCs (Experimental Procedures). The P_open_ was estimated to be 0.32 ± 0.11 for GCs and 0.34 ± 0.10 for SC boutons (Figure 2D, E), and the total number of VGCCs was 3.5-fold higher in GC boutons (GCs: mean = 236 ± 26, median: 184; SCs: mean = 65 ± 8, median: 48, p < 0.001, Mann-Whitney). Thus, the 3-fold difference in the volume corrected sAPCaTs between SC and GC boutons can be accounted for by differences in the total number of VGCCs per bouton.

To provide an independent estimate of the relative number of VGCCs, SDS-FRL of the Cav2.1 subunit of VGCCs was performed in SC and GC boutons. Specificity of Cav2.1 labelling was validated in cerebellar tissue samples from Cav2.1 cKO mice (Figure S2E-M). Saturating concentrations of a Cav2.1 antagonist (1 μM ω-agatoxin) completely blocked sAPCaTs in SC (97 ± 4 %, n = 7; Figure 2G, H) but only 52.4 ± 7.7 % (n = 8) in GCs, indicating that the Cav2.1 labeling represents half the VGCCs in GCs, but all channels in SCs. In agreement Cav2.2 labelling was also found in GC AZ (Figure S2N, O). The mean number of gold particles labelling the Cav2.1 subunit per bouton was 1.5 higher in GCs than in SCs (GC: 38 ± 19, median: 34, CV: 0.5, n = 426 boutons; SC: 26 ± 17, median: 22, CV: 0.65, n = 184 boutons; Figure 2I, J). Taking into account that Cav2.1 represents the total population of VGCCs in SC and only half in GC the 1.5x larger number of particles labelling Cav2.1 subunit actually indicates a 3-fold larger number of VGCCs in GCs than SCs; a value similar to that obtained from fluctuation analysis (Figure 2F). Taken together, our results suggest that differences in the total number of VGCCs is the principal factor responsible for the different sAPCaT amplitudes.

### Differences in VGCCs-Ca^2+^ sensor coupling strength

How might a 3-fold larger number of VGCCs in GC boutons be consistent with a lower P_v_ ? We next considered the possibility that VGCCs are more efficiently coupled to SVs in SC, possibly through a closer physical distance. In order to estimate the efficiency of VGCC to SV coupling we measured the fractional reduction in synaptic transmission before and after competition with the slow Ca^2+^ chelator, ethylene glycol-bis(β-aminoethyl ether)-N,N,N′,N′-tetraacetic acid (EGTA; Eggermann et al., 2012). GC-PC or GC-SC excitatory postsynaptic currents (EPSCs) were significantly more sensitive to a 20 min application of 100 μM EGTA-AM as compared to SC-SC IPSCs (GC-SC EPSC blockade: 51.1 ± 6.0 %, n = 10 cells; GC-PC EPSC blockade: 51.3 ± 5.0 %, n = 7 cells; SC-SC IPSC blockade: 18.4 ± 7.8 %, n = 10 cells; Figure 3A-C).

**Figure 3:**
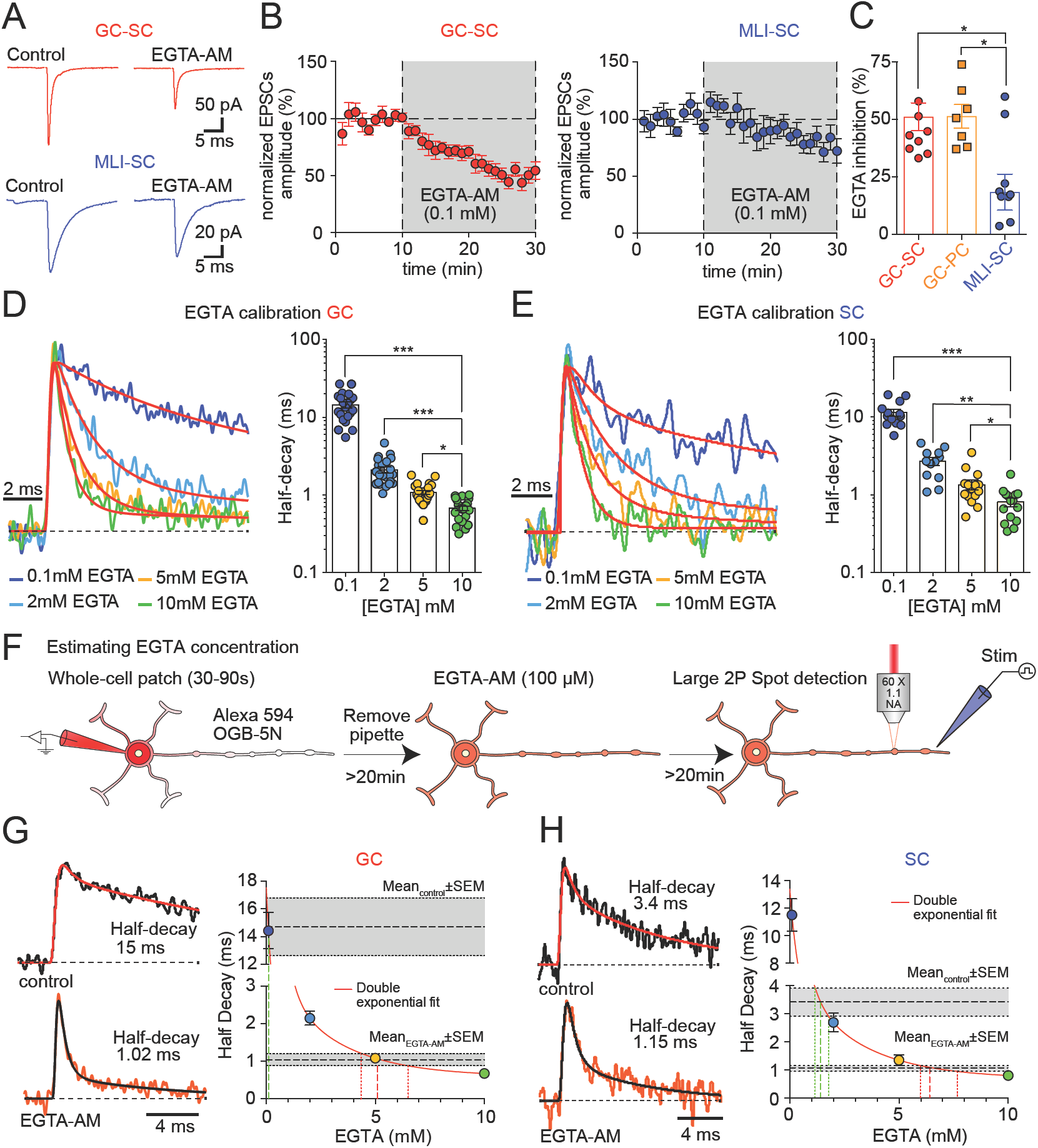
PF-EPSCs are more sensitive to EGTA inhibition than SC-IPSCs. **(A)** Representative traces of GC to Pukinje cell (red) and Molecular layer interneurons (MLI)-SC (blue) synaptic currents recorded before and after bath application of 0.1 mM EGTA-AM (right) for 20 minutes (min). Traces are averages of 60 extracellular stimuli delivered at 0.2 Hz before and 15 min after EGTA-AM addition. **(B)** Time course of EGTA-AM effect PF-evoked EPSCs (n = 10 cells, left) and SC-evoked IPSCs (n = 10, right). Error bars represent ± SEM. **(C)** EGTA-AM inhibition of synaptic currents after 20 min (average over 5 min) recorded from GC-PC (orange, n = 8), GC-SCs (red, n = 10), IPSCs recorded in SCs (blue, n = 10). (* p < 0.05, Kuskal-Wallis followed by Dunn’s multiple comparisons test) **(D)** *Left:* Normalized population averaged sAPCaTs recorded from GC boutons using large spot detection and OGB-5N (100 μM) in the presence of different intracellular concentrations of EGTA (0.1, 2, 5 and 10 mM). *Right*: Summary plot of the half decay times for all boutons. (* p < 0.05, **< 0.01, ***p< 0.001 Kuskal-Wallis followed by Dunn’s multiple comparisons test). **(E)** Same as D but for SC. **(F)** Schematic Illustration of experimental approach used to probe intracellular EGTA concentration in GCs boutons after bath application of EGTA-AM. **(G)** Population averaged sAPCaTs recorded using OGB-5N and large spot detection in GC boutons before (top left) and after bath application of EGTA-AM (0.1 mM; bottom left). Mean half-decay of control and EGTA-AM sAPCaTs (dashed line with grey region representing SEM) plotted on a calibration graph of sAPCaT half-decay with known [EGTA] (from panel C). Fine dashed lines define the estimated range of intracellular [EGTA], defined by the intersection of the SEM limits and the double exponential fit. **(H)** Same as in **G**, but for SCs

To exclude the possibility that the differential effect of EGTA-AM on synaptic transmission between GC and SC boutons results from differential intracellular accumulation of EGTA, we calibrated the intracellular concentration of EGTA using the decay of sAPCaTs recorded with OGB-5N. Neurons were whole-cell patch-clamped and dialysed for >20 min with an internal solution containing 100 μM of OGB-5N and different concentrations of EGTA (0.1, 2, 5 and 10 mM). Increasing intracellular EGTA concentrations decreased the half-decay of sAPCaTs recorded with two-photon (2P) large spot detection (Figure 3D-E). This fluorescence decay calibration curve was used to estimate the concentration of hydrolyzed EGTA in the previous experiments (Figure 3F). SC and GCs were next preloaded with OGB-5N, and after a 20 min bath application of 100 μM EGTA-AM the decay of sAPCaTs was accelerated in both SC and GC boutons (half-decay, SC: 1.1 ± 0.1 ms, n = 13; GC: 1.0 ± 0.2 ms, n = 11; Figure 3G, H). EGTA-AM produced a sAPCaT with a decay similar to that of ∼5 mM EGTA for GCs boutons and 6.5 mM in SCs (Figure 3G and H). Dialysis of SCs significantly slowed the decay of sAPCaTs, consistent with an endogenous buffer equivalent to 1.5 mM EGTA. Thus for SCs the added exogenous buffer also corresponded to an intracellular EGTA concentration of 5 mM. Therefore, the higher sensitivity of EPSC amplitudes at GC synapses to EGTA-AM application suggests a longer distance between VGCC and SV in GC boutons, which could also account for its lower P_v_.

### Similar duration of Ca^2+^ entry in GC and SC boutons

The duration of presynaptic AP has been shown to vary between synapses (Rowan et al., 2016) increases the duration of single VGCC openings (Borst and Sakmann, 1998; Sabatini and Regehr, 1997), and in turn can affect the interpretation of Ca^2+^ chelator experiments (Nakamura et al., 2015; 2018). But measuring AP duration in small synaptic boutons using classical electrophysiological techniques is challenging. We therefore used the first time derivative of sAPCaTs as a proxy for the duration of presynaptic Ca^2+^ currents (Sabatini and Regehr, 1998). sAPCaTs were recorded using 500 μM of OGB-5N with a large, stationary 2P excitation spot (Figure 4C, B). The derivative of the population averaged sAPCaT was fit with a Gaussian function, resulting in a full width at half maximum (FWHM) of 254 μs for GCs and 236 μs for SCs (Figure 4A-D). Moreover, we did not detect a difference in the rise times of sAPCaTs recorded from GC and SC boutons (GCs: 0.32 ± 0.03 ms, n = 41 boutons, 9 cells; SCs: 0.33 ± 0.03 ms, n = 41 boutons, 17 cells; p = 0.78 Mann-Whitney; Figure 4E). Taken together these results suggest that the duration of Ca^2+^ entry is similar between GC and SCs boutons and therefore can neither account for the difference in sAPCaT amplitude, and is consistent with the similarity in P_open_ from variance analysis, nor alter our estimation of VGCC-SV coupling distance as derived from the EGTA inhibition experiments.

**Figure 4:**
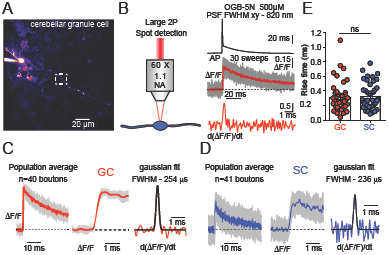
Ca^2+^ entry duration in GC and SC boutons is similar. **(A)** 2PLSM MPI image patch-labelled GC (Alexa-594) from which large spot 2P sAPCaTs were recorded. **(B)** Somatic AP (*top*) and its evoked sAPCaTs (*middle*) recorded using high cencentrations of OGB-5N (500 μM) from bouton in A. First derivative of the averaged (30 sweeps) ΔF/F trace (bottom). **(C)**: Population average (left) and expanded time scale (*middle*) of GC sAPCaTs (large spot detection using 500 μM OGB-5N). Grey is SEM. *Right:* First derivative (red trace) and Gaussian fit (black trace) of population averaged sAPCaTs. **(D)** Same as C but for SCs (n=72 boutons). **(E)** 10-90% rise times of sAPCaTs from C and D (n=41 GC boutons and n=71 SC boutons; p>0.05 Mann-Whitney test).

### Nanoscale distribution of VGCCs in GC and SC boutons

The differential sensitivity of synaptic currents to EGTA suggests that the distance between SVs and VGCCs is shorter in SC boutons despite the large number of VGCCs at AZs in the lower P_v_ GC boutons. We hypothesize that different VGCC-SV topographies are essential to account for the two findings we next examined the spatial patterns of gold particle labeling the Cav2.1 subunit in our SDS-FRL EM images (Figure 5A-B). We measured mean nearest-neighbor distances (NND) between gold particles and examined whether the observed NND distributions significantly differed from random arrangements (null model; for details see Experimental Procedures). Because the AZs in SC boutons cannot be delineated based on morphological features, we analyzed the gold particle distributions over the entire fractured surface of each SC bouton. In GC boutons the high density of intra-membrane particles allowed delineation of AZs and we analyzed the gold particle distributions only in the AZs of GCs (experimental procedures). For both SC and GC boutons the mean NND distributions of gold particles were shorter than expected from random arrangements (Figure 5A, B; p < 0.001, Mann-Whitney).

**Figure 5:**
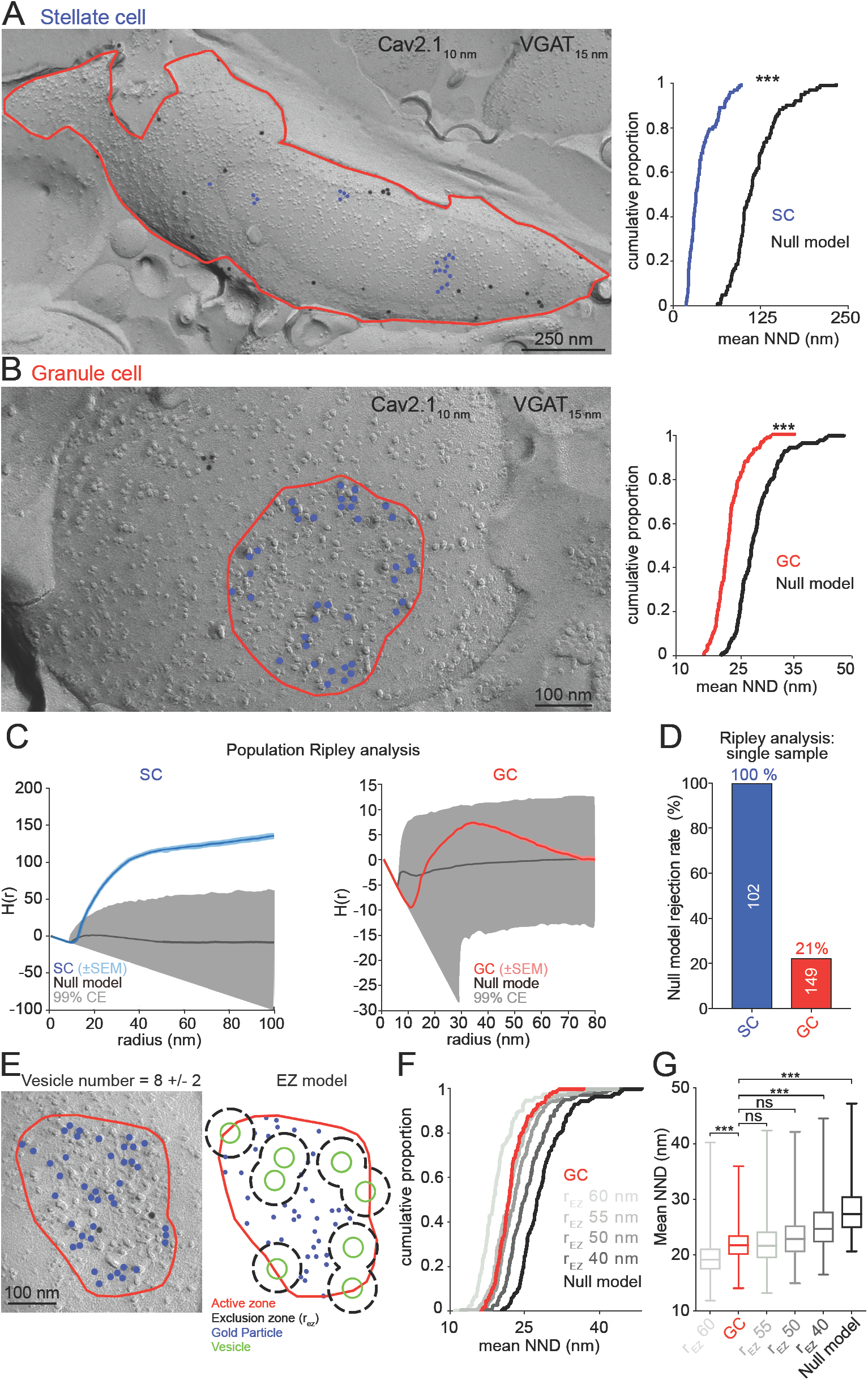
Nanoscale gold particle distribution in GC and SC boutons. **(A)** *Left*. High-resolution immunogold localization of the Cav2.1 subunit (blue dots) in a putative SC bouton delineated by the red line. *Right*. Cumulative distribution of mean NNDs (per AZ) of gold particles in SC boutons (n = 102 AZs; blue), and those of a null model (randomly distributed particles within bouton surface; generated from n = 1000 AZs, black). (p<0.001, Mann-Whitney test). **(B)** *Left*. High-resolution immunogold localization of the Cav2.1 (blue dots) subunit in a putative GC active zone (AZ, delineated by the red line). *Right*. Cumulative distribution of mean NNDs of gold particles in GC AZs (n = 149; red), and those of random distributions (n=1000) generated under the null model (black). The mean NND of the GC was significantly different from the null model (p<0.001, Mann-Whitney test). **(C)** Ripley H-function analysis of gold particle distributions across the population of SC (left) bouton surface and GC AZs (right). Shaded region indicates the 99% confidence envelope obtained from 5000 simulations of the null model for each SC bouton (red, n = 149) and GC AZ (blue, n = 102). The solid black line is the H(r) function for the null model. **(D)** Individual H-function analysis each bouton revealed that the null model was rejected (MAD test) for 100% of stellate cell (blue) and 21% for granule cell (red) patterns. **(E)** *Left*. Example electron micrograph of a putative GC AZ with Cav2.1 particles and a schematic (*right*) of a single gold particle pattern generated by the exclusion zone model (EZM) using the same number of particles and AZ area. **(F)** Cumulative distributions of mean NNDs for gold particle patterns in GC AZs (from B), EZM generated patterns (n = 1000 AZs for each exclusion radius, r_E_) and null model patterns (n=1000). **(G)** Box plot summary of mean NND of the data and EZM with different exclusion radii. (*** p < 0.001, Kuskal-Wallis followed by Bonferroni multiple comparisons test)

Previous analysis of gold particle labelling of Cav2.1 channels in CNS synapses suggested that VGCCs form clusters (Eltes et al., 2017; Holderith et al., 2012, Nakamura et al., 2015; Miki et al., 2017). We therefore performed a spatial point pattern analysis that test for deviations in the density of particles over various spatial scales, using the Ripley’s H-function, an analysis method that is particularly sensitive to particle clustering (down to two points per cluster, for observed NNDs; Figure S3A-C). A Ripley’s H-function analysis of the entire population of SC boutons revealed a clustered gold particle distribution (p < 0.05, MAD test, Figure 5C left), whereas in GC AZs, the distribution was not significantly different from random (p = 0.315, MAD test, Figure 5C right). As the variability in AZ size and gold particle number might contribute to increased variance across the population precluding detection of clusters, Ripley analysis was performed on individual boutons and AZs. 100% of SCs boutons showed statistically significant deviation from the null model generated for each AZ, whereas only 21% was different in GC AZs (Figure 5D). Thus for SC boutons, the mean NND and H-function analysis are compatible with clustered models of VGCC distributions. For GC boutons, however, the H-function analysis suggested that Cav2.1 gold particle distributions were not random, but were *not* consistent with a cluster model.

Cluster analysis using the DBSCAN algorithm (see Supplementary Information, Figure S3D-K) of gold particle distribution in SC indicated that each SC bouton comprised an average of 4.7 ± 2.7 clusters (mean ± SD; DBSCAN parameters: ε = 50 nm, minimum number of gold particles per cluster = 2; Figure S3G). The number of gold particles per cluster was of 5.2 ± 3.5 (mean ± SD; n = 102 boutons; Figure S3H). The mean area of single clusters was 10-fold smaller than the total estimated synaptic area (Figure S3I; Nusser et al., 1997), as was observed in the calyx of Held where a PCM model is thought to drive release. The intercluster distance is too long (median of 245 nm, Figure S3J) to cooperatively contribute to SV fusion (> 100nm distances drive undetectable levels of release, Nakamura et al. 2015). Thus together these data are compatible with the presence of multiple small VGCCs clusters per SC bouton, each potentially driving release using a PCM topography.

Previous modeling studies have proposed VGCC-SV topographies in which VGCCs are excluded from a zone around synaptic vesicles (EZM; Keller et al., 2015). We considered whether such a model could account for the particle distribution with single GC AZs. We therefore generated point patterns by first placing multiple docked vesicles (8 ± 2; Xu-Friedman et al., 2001) randomly within the AZ (Experimental Procedures), then placing VGCCs (same number as obtained experimentally) within the AZ, but not closer than a minimum distance from the center of a vesicle (exclusion radius, r_ez_; Figure 5E). When we compared the distributions of mean NNDs obtained from the model simulation and that of the experimental data, we found that an exclusion radius of 50 - 55 nm was compatible with the data (p < 0.05, Kruskal-Wallis test; Figure 5F, G). H-function analysis of generated particle distributions using EZM (50 nm radius) revealed no difference from the null model at the population level and a rejection rate of 32% for individual AZ, similar to the 21% rejection rate obtained for GC gold particle distributions (Figure S4A-F). The similarity between data and model generated point patterns, confirms that VGCCs do not form clusters at the GC AZs, and their distribution is consistent with a simple EZM in which the exclusion radius is 50 - 55 nm. If we reduced the number of docked vesicles (down to 4 ± 1) a slightly larger exclusion radius (60 - 75 nm) was needed to match the experimental NND distribution (p < 0.05, Kruskal-Wallis test; Figure S4G-I). Taken together, the spatial point pattern analysis method suggests that differences in the VGCC distributions at the two synapse populations can be accounted by two contrasting models: a clustered model for SC boutons and an EZM for GC AZs.

### Different nanoscale distributions of VGCCs account for functional diversity

To test whether cluster and EZM VGCC distributions could reproduce the different functional properties of GC and SC synapses, we implemented particle-based Monte Carlo simulations of Ca^2+^ reaction, diffusion and binding to a release sensor to predict P_v_ and EGTA inhibition of the postsynaptic responses. All model parameters were constrained by experimental measurements (Table S1), except the location of the Ca^2+^ sensor for release. Since SC channel distributions at the bouton surface were found to be clustered (as suggested in Figure 5), we performed simulations using perimeter release model (PRM) arrangement of channels and release sensors (Nakamura et al., 2015). P_v_ and EGTA inhibition were simulated for a release sensor located at various distances from the edge of a cluster of VGCCs containing either 9, 16 or 32 VGCCs corresponding to the presence of either 7, 4 or 2 of clusters of VGCCs per bouton (Figure 6A, Figure S3). The AP waveform driving release was adjusted to produce a Ca^2+^ entry time course and VGCC P_open_ (Figure S5) matching experimental observations (Figure 4 and 2, respectively). The predicted EGTA inhibition was compatible with our experiments (19 ± 8%, n = 10 cells; Figure 4A-C) if the coupling distance was < 30 nm (Figure 6B). Simulated P_v_ was highest when the Ca^2+^ sensor was placed 10 nm from the cluster’s edge, and ranged from 0.27 with 9 channels per cluster up to 0.5 for 32 channels per cluster in agreement with the range of published values (P_v_ = 0.3 - 0.8; Arai and Jonas, 2014; Pulido et al., 2015; Figure 6C). This simulation supports a PRM in which clusters of VGCCs drive the fusion of SVs that are tightly coupled to cluster perimeters (10 - 20 nm).

**Figure 6:**
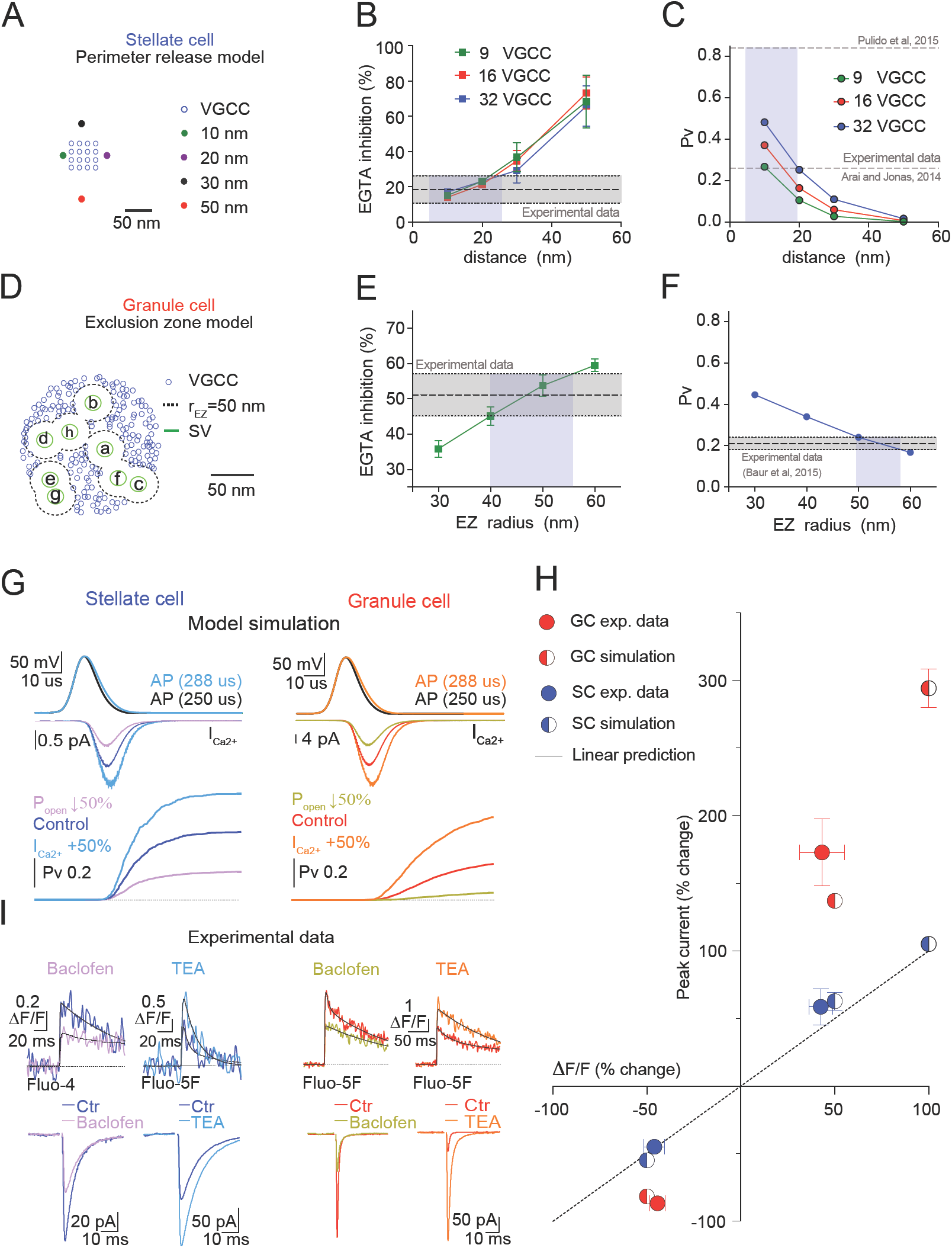
Stochastic simulations of vesicular release from PRM and EZM reproduce SC and GC synaptic transmission. **(A)** VGCC distribution representing that of SC clusters used for Monte Carlo (MC) simulations of Ca^2+^ entry, reaction-diffusion, and binding to sensor for vesicle fusion. 9, 16 (example shown) and 32 channels per cluster with a mean channel NND of 15 nm were tested. The Ca^2+^ sensor was placed 10, 20, 30 and 50 nm from the nearest perimeter channel (perimeter release model, PRM). **(B, C)** Simulated 5 mM EGTA-inhibition (**B**) and P_v_ (**C**). Symbols represent mean±SEM of 5000 simulation trials. Grey zone in B represents the mean±SEM of experimental EGTA-inhibition of SC-SC synapses (Fig. 3). Grey zone in C indicate published P_v_ values (Arai and Jonas, 2014; Pulido et al., 2015). Light blue boxes indicate simulated coupling distances compatible with experiments. **(D)** EZ arrangement of VGCC used to probe vesicle release probability in GCs. Ca^2+^ sensors were placed at center of vesicle (green), which is also the center of the EZ. **(E-F)** Simulated 5 mM EGTA-inhibition (**E**) and P_v_ (**F**) for EZM VGCC distributions generated with indicated r_E_. Symbols represent mean±SEM of 5000 simulation trials. Gray zones in E indicates the mean±SEM of experimental EGTA-inhibition of GC-PC synapses (Fig. 3). Grey zone in F indicates mean±SEM of published P_v_ (Baur et al., 2015). Light blue boxes indicate simulated coupling distances compatible with experiments. **(G)** AP waveforms (top) and MC simulated calcium currents (middle) and cumulative release plots (bottom) using either PRM (SC) or EZM (r_E_ = 50 nm, GC). P_v_ was estimated from simulations in which Ca^2+^ currents were either increased (SC: light blue, n = 1000; GC: orange, n = 1000) by broadening the AP, or decreased (SC: pink, n = 5000; GC: lime, n = 2000) by 50% compared to control (SC: blue, n = 2000; GC: red, n = 1000) by reducing open probability of VGCCs. **(H)** Experimental Fluo-5F sAPCaTs (200 μM; top) recorded before and after application of Baclofen (10 μM) or TEA (GC: 300 μM, SC: 1 mM). TEA increased peak sAPCaTs in GCs (43.3 ± 12.1%, n = 9 boutons, 4 cells; red) and SCs (42.6 ± 6.3%, n = 10 boutons, 4 cells; blue). *Bottom:* Representative GC-PC EPSCs (red) or SC-SC IPSCs (blue) before and after Baclofen or TEA application. Increase PSC TEA: SCs: 58.6 ± 13.6%, n = 8 cells; GCs: 173.9 ± 25.0%, n = 8 cells. **(I)** Modulation of sAPCaT against modulation of synaptic currents (EPSCs or IPSCs) by TEA and Baclofen. The dashed line is a slope of 1.

For GC synapses we performed particle-based simulations using our EZM (Figure 5D) to distribute VGCC within the AZ. A Ca^2+^ sensor was placed at the center of exclusion zones with various exclusion radii (30 - 60 nm; Figure 6D, E). An exclusion radius of 50 nm predicted a P_v_ (0.24 ± 0.005; Figure 6F) and an EGTA inhibition (53.8 ± 3.0%; Figure 6F) that closely matches the experimental obtained values (Figure 4). Different distributions of SVs had only slight effect on simulated EGTA inhibition and P_v_ values (Figure S6M), indicating that the specific arrangement of SV is not a major determinant of EGTA inhibition and P_v_ values in EZM.

A longer coupling distance together with higher number of VGCCs present in GC boutons could manifest a higher cooperativity between VGCC activation and release (Matveev et al., 2011), effectively enhancing the sensitivity of release to neuromodulation. In order to test our hypothesis, we used Monte Carlo simulations to compute P_v_ using either PRM (10 nm) for SCs or EZM (50 nm) for GCs and evaluated the impact of altering the activity of VGCCs to mimic the action of neuromodulators by increasing or decreasing VGCC P_open_ (Figure 6G). Our simulations revealed that for the SC PRM, altering P_open_ translated into a linear modulation of P_v_ (Figure 6G, H), consistent with only a few VGCCs driving release (Augustine, 1990; Eggermann et al., 2012; Luo et al., 2015). In contrast, simulations using EZM (50 nm) predicted a nonlinear reduction in P_v_ of 82 ± 2% for a 50% decrease in Ca^2+^entry and a nonlinear increase of 137 ± 5% for a 50% increase in Ca^2+^entry (Figure 6G, H) supporting multiple channel driving release (Dittman and Regehr, 1996; Mintz et al., 1995).

To test these model predictions, we experimentally manipulated presynaptic Ca^2+^ entry in response to single APs in both SC and GC boutons and measured the corresponding effects in sAPCaTs and neurotransmitter release through electrophysiological recording of postsynaptic currents. The GABA_B_ receptor agonist, baclofen (10 μM) reduced the amplitude of sAPCaTs to a similar extent in both SC and GC boutons (Figure 6H; GC: 44.3 ± 4.2 %, n = 7 boutons, 3 cells; SC: 46.0 ± 5.6 %, n = 10 boutons, 3 cells). In agreement with model predictions, despite similar reduction in the amplitude of sAPCaTs, baclofen application produced a significantly larger decrease in synaptic strength in GC as compared to SCs synapses (Figure 6I-H; SCs: 44.8 ± 4.6%, n = 9 cells; GCs: 86.6 ± 4.5%, n = 4 cells, p = 0.01, Mann Whitney test). Model predictions were also confirmed when increasing the Ca^2+^ entry using the K^+^ channel blocker TEA (1 mM SCs and 300 μM GCs; p = 0.003, Mann Whitney test, Figure 6I, H). In conclusion, the differential cooperativity between VGCCs and SVs at SC and GC synapses is consistent with the different nanotopographical arrangements of VGCCs and SVs, and tune the sensitivity of neurotransmitter release to the action of neuromodulators.

Thus the data from SC boutons are most consistent with a VGCC-SV topography compatible with a PRM as in the Calyx of Held (Nakamura et al., 2015). In contrast, both functional and nano-anatomical VGCC data from GC boutons are consistent with an unconventional VGCC-SV topography in which VGCCs are excluded from a 50 nm zone around the Ca^2+^ sensor.

### Munc13-1 labelling confirms novel nanotopographical arrangement of VGCC and SVs

In order to more directly test the EZM model by direct localization of VGCCs and putative SVs, we performed immunogold co-localization of Cav2.1 and Munc13-1 molecules (Figure 7A) in the same preparations. It has recently been reported that Munc13-1 is a core component of the vesicle release site (Sakamoto et al., 2018). We therefore used Munc13-1 as a molecular indicator of the vesicle docking site. In contrast to Cav2.1 gold particles (Figure 5), Ripley’s H-function analysis of Munc13-1 gold particle distributions in GC AZs showed significant deviation from the null model (P < 0.05) in a manner compatible with clustering of Munc13-1 molecules (Figure S6). Cluster analysis revealed 7.1 ± 3.4 Munc13-1 clusters per GC AZ in agreement with previous EM estimates of the number of docked SVs (Xu-Friedman et al., 2001) that also scaled linearly with AZ area (Figure S6). These results thus suggest that Munc13-1 clusters are a good marker for identifying putative release sites.

**Figure 7:**
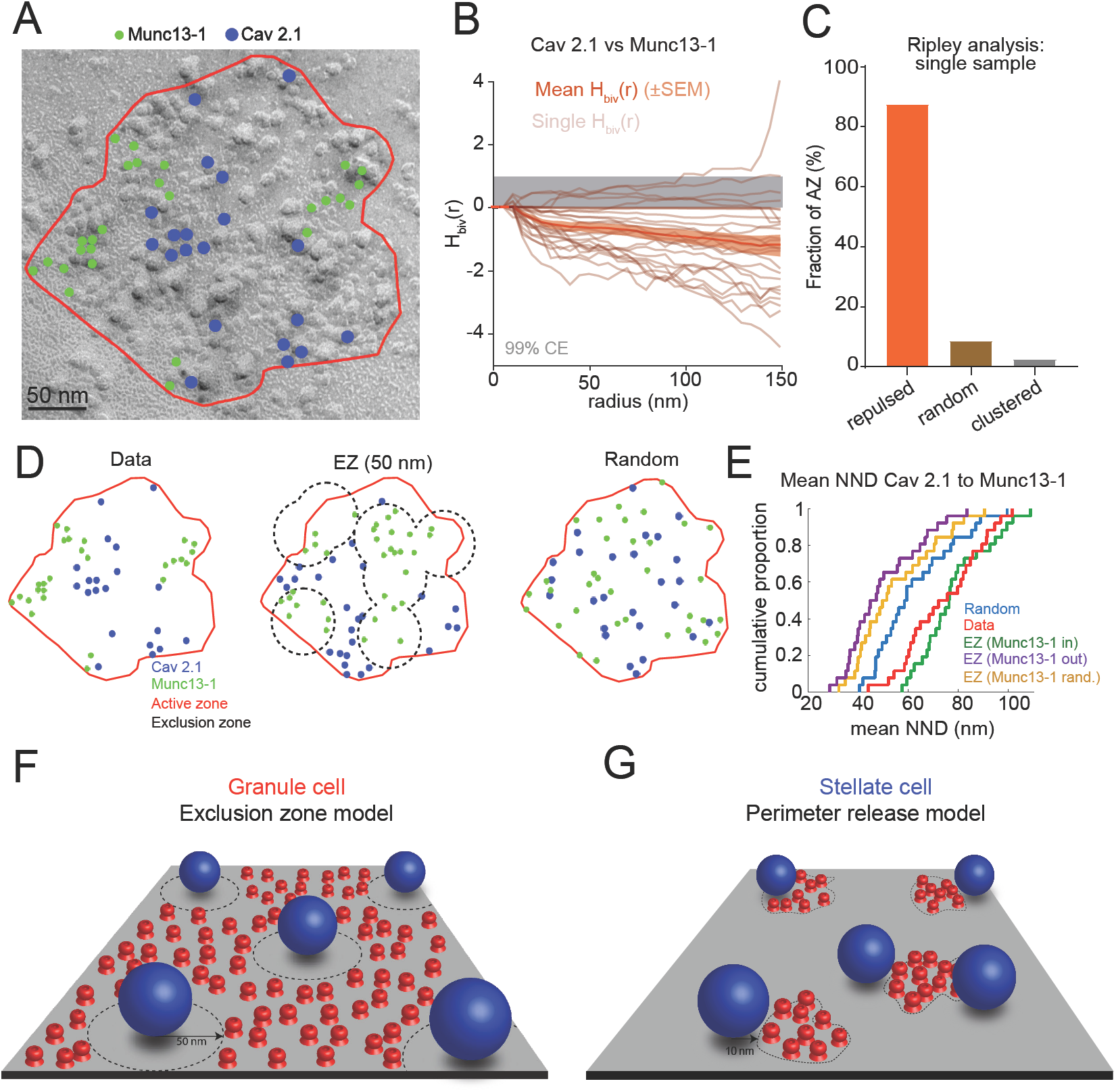
Nanotopographical arrangement of VGCCs and Munc13-1 are consistent with EZM for GC synapses. **A**- High-resolution immunogold localization of Cav2.1 subunit (blue) and Munc13-1 (green) in a GC AZ. **B**- Ripley bivariate H-function analysis of Munc13-1 and Cav2.1 gold particle spatial patterns across 26 GC AZs indicates two par. Shaded region indicates the 99% confidence envelope obtained from 500 simulations of the null model for each GC AZ. Light orange curves represent the normalized Ripley bivariate function (see Methods). Dark orange curve represents population mean ± SEM (n = 26). **C**- Percentage of GC AZs in which the bivariate H-function suggested random (3%), clustered (9%) or repulsed (88%) interaction between Cav2.1 and Munc13-1 point patterns. Paired analyses were performed using the same area and number of each particle for each AZ and the null model (500 simulated spatial patterns per AZ area). Null model rejection was determined using a MAD test. **D**- Representative single Munc13-1 and Cav2.1 gold particle patterns obtained from the electron micrograph of GC AZ shown in G (left) as well as patterns generated using EZM (r_E_ = 50 nm) VGCC distribution with Munc13-1 particles placed randomly within EZ (middle) and a simulated null model for both particles within the AZ (right). **E**- Cumulative mean NND distribution between Munc13-1 and Cav2.1 gold particles experimental data (red), EZM models in which Munc13-1 was placed inside (red), outside (blue) EZs, or randomly placed within AZ (yellow). 100 patterns were generated for each AZ. **F-G**- Summary illustration of nanotopographical arrangement of VGCCs (red) and SVs (blue spheres), suggesting GCs use an EZM and SCs use a PRM.

To analyze the relative spatial distributions of gold particles labeling Cav2.1 and Munc13-1, we used a bivariate extension of Ripley’s H-analysis (H_biv_(r)) that provides information about spatial correlation between two different point patterns (Hanisch, 1979). Bivariate analysis of 26 GC AZs revealed a negative correlation between Cav2.1 and Munc13.1 gold particles in GC AZs that was statistically different (p < 0.05) from that generated by a null model in both population and single AZ analysis. Such analyses suggest an apparent repulsive interaction between Cav2.1 and Munc13.1 gold particles (Figure 7B, C), or at least a mechanism that ensures the two proteins are not close to each other. Such an anti-correlation would be expected if Munc13-1 clusters were located within the EZ proposed for VGCC distributions.

To test if an EZM could account for both the Munc13-1 and Cav2.1 distributions, we simulated particle distributions in individual AZs using different arrangements of the two proteins and compared the resulting Cav2.1 to Munc13-1 NNDs to that of experimental data. For each AZ we added 8 ± 2 docked vesicles (scaled by AZ size), and placed Cav2.1 particles according to EZM (exclusion radius: 50 nm). Munc13-1 particles were either placed within (EZ_munc13-1in_) or outside EZs (EZ_munc13-1out_) or randomly distributed within the AZ (EZ_munc13-1random_; Figure 7D, E). We also tested the condition where both Cav2.1 and Munc13-1 particles were randomly distributed in the AZ. The experimental Cav2.1 to Munc13-1 mean NND distribution was statistically different from all groups except for the case where Munc13-1 was placed within the EZ (Figure D, E). In summary, the nanotopographical arrangement of Cav2.1 and Munc13-1 particles is described by an EZM where SVs are docked within a sea of randomly placed VGCCs within an AZ, but with a minimum distance of 50 nm from the closest VGCC.

All the data taken together, the functional and structural data revealed that GC and SC boutons use two strikingly different VGCC-SV nanotopographical motifs (Figure 7F, G) to achieve different functional behaviors.

## Discussion

While many of the molecules involved in orchestrating signaling cascades driving submillisecond synaptic transmission are known, how the nanoscale topography of the major molecular players could be tuned to manifest in the heterogeneous function at small bouton-type CNS synapses is not well understood. We provide here the first nanoscale view of how the molecular organization of VGCCs and SVs governs synaptic strength and sensitivity to neuromodulation. We showed that in strong SC synapses SVs are tightly associated with the perimeter of VGCC clusters, while for the weaker GC synapses VGCCs are displaced away from putative docked SVs. These strikingly different topographical motifs provide novel insights into how specific molecular topographies can contribute to functional heterogeneity at synapse, and put into question the simple notion that bouton Ca^2+^ entry can be used as a proxy for P_v_ (Holderith et al., 2012; Koester and Johnston, 2005; Nakamura et al., 2015). Also unexpected was the observation that VGCC organization does not necessarily follow the same clustering rules as for aligning SVs and postsynaptic ionotropic receptors (Biederer et al., 2017; Hruska et al., 2018; Sakamoto et al., 2018; Tang et al., 2016). Finally, the two topographical models (PRM and EZM) point to different macromolecular organization within the presynaptic terminal; one is responsible for clustering channels and tethering SVs, while the other strategy requires a putative molecular spacer.

### Deriving function from nanoscale structure

Recent work at the calyx of Held and subsequent simulations indicate that if the nanoscale distribution of VGCC is known, then the experimental observations such as EGTA inhibition of neurotransmission, P_v_ and release time course, can be used to constrain VGCC-SV topographies (Chen et al., 2015; Nakamura et al., 2015). However, these methods require extensive knowledge of the Ca^2+^ entry diving release (Nakamura et al., 2018), and thus have been limited to terminals accessible by patch-clamp methods. We circumvented this limitation by employing a derivative method (Sabatini and Regehr, 1998) to estimate the time course of Ca^2+^ entry, and an fluctuation analysis method (Sabatini and Svoboda, 2000) to estimate P_open_ and the total number of VGCCs within single, small presynaptic boutons. While only the total number of VGCCs differed between strong and weak synapses, estimating their P_open_ and the Ca^2+^ entry time course were also necessary for accurate simulations of P_v_ and EGTA inhibition (Nakamura et al., 2015; 2018).

The precise localization of VGCCs is an essential measurement for inferring the nanoscale synapse topography of key molecules. Our results revealed a clustered distribution in SC boutons, and a non-random, non-clustered distribution in GC boutons that is consistent with an EZM. These VGCC distributions were then used to create particle-based simulations of Ca^2+^ reaction and diffusion. EGTA inhibition, P_v_ and neuromodulation were well predicted by 10 nm (but not 20 nm, Figure S7) coupling distances for SCs and 50-55 nm distances, which matched the EZ radius that predicted the Cav2.1 gold particle distributions in GCs. Using Munc13-1 as a marker for release sites (Sakamoto et al., 2018), we analyzed for the first time the relative distribution of Cav2.1 and the SV docking sites in single GC AZs. In contrast to Cav2.1, Munc13-1 gold particles were clustered and spatially anticorrelated with VGCCs, a finding consistent with the EZM.

Thus a combination of single bouton optophysiology, nanoscale molecule localization analysis and Monte Carlo simulations, provided unique insights into not only VGCC-SV distances, but also their nanoscale topographical arrangement.

### Putative molecular players generating topographical motifs

VGCC-SV topographies are likely dictated by the organization of synaptic proteins into macromolecular complexes, for which Rab3-interacting molecules (RIMs), RIM binding proteins (RPBs), and Munc13 proteins are likely to be key players (Han et al., 2011; Kaeser et al., 2011). Loss of all RIM proteins was shown to impair fast neurotransmitter release from tightly coupled SVs in hippocampal cultures and at the calyx of Held. Because the calyx of Held is a tightly coupled synapse that uses a PRM topography, it is conceivable that the presence or absence of RIM proteins may define PRM or EZM topographies, respectively. However, a simple lack of tethering might not reproduce the rather specific and long (50 nm) minimal distance between SVs and the closest VGCC that we found at GC synapses.

Munc13-1 is thought to couple SVs and VGCCs via Rab3 and RIM proteins (Deng et al., 2011). However, there is also evidence in cultured neurons that Munc13-1 directly interacts with VGCCs (Calloway et al., 2015). In Drosophila NMJs, the N-terminal domain of the long Unc13A isoform is capable of tethering vesicles at a particular distance (∼70 nm from VGCC cluster center; Reddy-Alla et al., 2017), which implicates that these large proteins tether at specific distances. Indeed, the Unc13B isoform, which has a larger N-terminal domain, is itself localized at longer distances (120 nm from VGCC cluster center), and is responsible for tethering vesicles at a similar distance (Bohme et al., 2016). In rodent synaptic terminals there are four isoforms Munc13-1, uMunc13-2, ubMunc13-2 and Munc13-3. Munc13-1 is expressed at the majority of synapses, whereas the others can be synapse-specific. In hippocampal mossy fibers, loss of Munc13-2 leads to a reduction of release probability (Breustedt et al., 2010). Because hippocampal MFs exhibit loose VGCC-sensor coupling (>60 nm; Vyleta and Jonas, 2014), it is interesting to consider the possibility that Munc13-2 might be specific to synapses exhibiting longer VGCC-sensor coupling distances. Munc13-1 and -3 are expressed in cerebellar GC synapses (Augustin et al., 2001), which we have shown here to exhibit long coupling distances. While Munc13-3 (the largest Munc13 protein) is essential for normal release at these terminals (Kusch et al., 2018), it could also contribute to the longer coupling distances we observed in GCs (> 50 nm). Finally, in hippocampal neurons, loss of RIM-BP2 is responsible for increased VGCC-sensor distances, and represents another candidate molecule potentially influencing the nanoscale topography of VGCCs and SVs (Grauel et al., 2016).

### Physiological implications of different topographical motifs

One of the principal differences between the two motifs presented here is the physical distance between VGCCs and the Ca^2+^ sensor for SV fusion. Short coupling distances (< 20 nm) are0020associated with high P_v_ (Rozov et al., 2001), short synaptic delays (Bucurenciu et al., 2010; Nakamura et al., 2015) and a brief time course of release (Nakamura et al., 2015). Tight VGCC-sensor coupling could also account for an insensitivity of release time course to the magnitude of the Ca^2+^ influx (Arai and Jonas, 2014; Datyner and Gage, 1980; Sargent et al., 2005; Van der Kloot, 1988). Thus, tight coupling maintains temporally precise and reliably synaptic transmission despite alterations in Ca^2+^ influx via neuromodulators or through VGCC upregulation by homeostatic mechanisms (Glebov et al., 2017). SCs synapses exhibit all of these features (Arai and Jonas, 2014; Pulido et al., 2015), and thus are well poised to finely tune PC excitation through precise feedforward inhibition (Mittmann et al., 2005).

GC synapses, in contrast, exhibit loose VGCC-sensor coupling using an entirely different VGCC-SV topography that is consistent with an EZM. The unique topography with long coupling distances and many VGCCs within the AZ results in a higher P_v_ than expected for long (>50 nm) coupling distances (P_v_ < 0.1, Figure 6F; Nakamura et al., 2015; Vyleta and Jonas, 2014). As is the case with PRM, long coupling distances in the EZM (large EZ radii) also slow the time course of release (Figure S6K, L). The long coupling distance, also imparts a supralinear relationship between number of open channels and release, which is not the case for tight coupling where only few channels drive release (Fedchyshyn and Wang, 2005; Matveev et al., 2011). Thus vesicular release is less sensitive to the stochasticity of single channel openings, yet supralinearly sensitive to VGCC P_open_ during the AP. Neuromodulators are therefore likely to differentially modulate excitation and inhibition, potentially altering E/I balance in the molecular layer of the cerebellum. The large number of channels may also be important for generating a large residual [Ca^2+^] that can efficiently drive other Ca^2+^ dependent signaling cascades, in particular those that drive short-term facilitation (Jackman and Regehr, 2017; Turecek and Regehr, 2018).

The EZM also lends itself to a high degree of heterogeneity in P_v_ due to variable number of VGCCs within close proximity to the sensor (see different possible topographies, Figure S5). One advantage of release heterogeneity is the ability of single axonal connections to convey both transient changes in firing rate using high P_v_ vesicles, as well as for reporting tonic changes in firing rate using low P_v_ SVs. Thus single AZs with multiple release sites and heterogeneity in P_v_ seem to be particularly useful for faithfully transmitting the temporally rich information in GC firing rates (Chabrol et al., 2015). This is of particular importance if GCs make only one synaptic contact onto PCs (Isope and Barbour, 2002).

### Are the PRM and EZM canonical motifs?

PRM and EZM represent opposing nanoscale molecular topographies, which can mediate tight and loose coupling between VGCCs and SVs, respectively. To date some of the most well accepted nanoscale topographical motifs are those described for the vertebrate NMJ that are consistent with rows of calcium channels and SVs (Harlow et al., 2001; Robitaille et al., 1990). Recent numerical simulations of species-specific differences in those motifs (Nagwaney et al., 2009) could account for heterogeneity in P_v_ and STP (Laghaei et al., 2018). While these models are compelling, the precise nanoscale location of VGCCs has not been determined. In drosophila NMJ, super-resolution microscopy has demonstrated that VGCCs are arranged in a tight cluster (50 - 100 nm diameter) with vesicles being tethered around its perimeter (Liu et al., 2011). Recently at mouse inner hair cells super-resolution microscopy identified linear clusters of VGCCs (Neef et al., 2018; Wong et al., 2014) that in combination with VGCC cooperativity measurements and numerical simulations drive release from within ∼20 nm of the cluster perimeter (Pangrsic et al., 2015).

Structural evidence for distinct VGCC-SV topographies in CNS synapses is well described at the calyx of Held, where a PRM is thought to drive SV fusion (within ∼12-20 nm; Nakamura et al., 2015). Nevertheless, numerical simulations mimicking physiological results at other CNS bouton-type synapses proposed different topographies, such as clustered Ca^2+^ channels and random vesicle placement within the AZ of cultured hippocampal boutons (Ermolyuk et al., 2013) or random placement of both SVs and VGCCs (Scimemi and Diamond, 2012) at hippocampal Schaffer collateral boutons. Examination of these synapses with SDS-FRL in combination with biophysical characterization of Ca^2+^ flux, P_v_ and EGTA-inhibition measurement could provide insight into whether other nanoscale topographical motifs of VGCCs and SVs exist. Nevertheless, findings in the literature suggest that PRM topographies are used at synapses with tight VGCC-SV coupling (e.g. inner hair cells, calyx of Held synapses, cerebellar MFs (Delvendahl et al., 2015), fast-spiking interneurons (Bucurenciu et al., 2008) and SCs, while EZM topographies are use at synapses with loose coupling (GCs or hippocampal mossy fiber synapses (Vyleta and Jonas, 2014).

## Acknowledgements

This study was supported by the Centre National de la Recherche Scientifique, Fondation pour la Recherche Medicale, Agence Nationale de la Recherche (ANR-2010-BLANC-1411 and ANR-13-BSV4-0016), Ile de France (Domaine d’Intérêt Majeur MALINF). The laboratory of D.A.D. is a member of the BioPsy Laboratory of Excellence. We would like to thank Florian Rückerl for technical assistance and Jeremy Dittman and Angus Silver for critical reading of the manuscript.

## Methods

### Slice Preparation

All animal experimental procedures were approved by the ethics committee CEEA - Paris1. Acute parasagittal slices (200 μm) were prepared from adult CB6F1 mice (F1 cross of BalbC and C57Bl/6J), both males and females, between 30 to 61 days of age. Mice were killed by rapid decapitation, after which the brains were quickly removed and placed in an ice - cold solution containing (in mM): 2.5 KCl, 0.5 CaCl_2_, 4 MgCl_2_, 1.25 NaH_2_PO_4_, 24 NaHCO_3_, 25 glucose, 230 sucrose, and 0.5 ascorbic acid bubbled with 95% O_2_ and 5% CO_2_. Slices were cut from the dissected cerebellar vermis using a vibratome (Leica VT1200S). After preparation, the slices were incubated at 32°C for 30 minutes in the following solution (in mM): 85 NaCl, 2.5 KCl, 0.5 CaCl_2_, 4 MgCl_2_, 1.25 NaH_2_PO_4_, 24 NaHCO_3_, 25 glucose, 75 sucrose and 0.5 ascorbic acid. Slices were then transferred to an external recording solution containing (in mM): 125 NaCl, 2.5 KCl, 2 CaCl_2_, 1 MgCl_2_, 1.25 NaH_2_PO_4_, 25 NaHCO_3_, 25 glucose and 0.5 ascorbic acid, and maintained at room temperature for up to 6 hours.

### Electrophysiology

Whole-cell patch-clamp recordings were performed at near physiological temperatures (33-35°C) using a Multiclamp 700B amplifier (Molecular Devices), and using fire-polished thick-walled glass patch electrodes (1.5 mm OD, 0.75 mm ID, Sutter Instruments; 5–10 MΩ tip resistance for GC and 4-6 MΩ for SC).

For Ca^2+^ imaging experiments pipettes were backfilled with the following internal solutions (in mM): GC-110 K-MeSO_3_, 40 HEPES, 0.1 EGTA, 4 MgCl_2_, 0.02 CaCl_2_, 0.3 NaGTP, 4 NaATP, 10 K_2_ phosphocreatine (∼280 mOsm pH adjusted to 7.3 using KOH); SC-120 K-MeSO_3_, 40 HEPES, 0.1 EGTA, 4 MgCl_2_, 0.02 CaCl_2_, 0.3 NaGTP, 4 NaATP, 10 K_2_phosphocreatine (∼300 mOsm pH adjusted to 7.3 using KOH). Alexa 594 (10 μM) and the Ca^2+^-sensitive dye (OGB-5N or Fluo-5F) at the desired concentration were added to the intracellular solution daily. The membrane potential (Vm) was recorded in current clamp mode (Multiclamp700B amplifier) and held at −89 mV ± 2mV (GC; Chabrol et al., 2015) and −70 Mv ± 2mV (SC), if necessary, using current injection (typically <-10 pA for GC and <150-pA for SC). A liquid junction potential of 7 mV (calculated using JPCalcW) was used to correct all membrane voltage values. Series resistance was compensated online by balancing the bridge and compensating pipette capacitance. APs were initiated by brief current injection ranging from 200 to1000 pA and 0.5-1 ms duration.

For voltage-clamp recordings (Multiclamp700B amplifier), cells were patched using the following intracellular solution: 120 K-MeSO3, 40 HEPES, 1 EGTA, 4 MgCl2, 0.49 CaCl2, 0.3 NaGTP, 4 NaATP, 10 K2 phosphocreatine (∼300 mOsm pH adjusted to 7.3 using KOH). Extracellular voltage pulses (50 μs, 5-50 V; Digitimer Ltd, UK) were delivered using a second patch pipette filled with ACSF and placed over SC or PC soma to stimulate synaptic responses while minimizing dendritic sublinearities (Abrahamsson et al., 2012). Stimulation intensity was adjusted to obtain a stable paired-pulse ratio, as described in (Abrahamsson et al., 2012). Series resistances (SC:15 ± 4 MΩ, mean and s.d., n = 104, GC:29 ± 6 MΩ, mean and s.d., n = 70) were not corrected online for SCs. PC recordings were performed with patch series resistances between 4 and 10 MΩ followed by 85% compensation. Data were discarded if series resistance was >30 MΩ or changed by more than 20% across the course of an experiment. SR 95531 (2-(3-carboxypropyl)-3-amino-6-(4-methoxyphenyl)pyridazinium bromide, 10μM) was added to the ASCF to block GABA_A_ receptors. In some experiments Baclofen (10 μM) and TEA (0.3 or 1 mM) were used to active GABA_B_ receptors or block potassium channels respectively (all drugs were purchased from Abcam, Cambridge, UK). Alexa-Fluor 594, OGB-5N, Fluo-5F were purchased from Life Technologies, USA. All recordings were low-pass filtered at 10 kHz and digitized at 100 kHz using an analog-to-digital converter (model NI USB 6259, National Instruments, Austin, TX, USA) and acquired with Nclamp software (Rothman and Silver, 2018) running in Igor PRO (Wavemetrics, Lake Oswego, OR, USA).

### Cellular imaging

Cells were identified and whole-cell patch-clamped using infrared Dodt contrast (Luigs and Neumann, Ratingen, Germany) and a frame transfer CCD camera (Scion Corporation, Cairn Research Ltd, Faversham, UK). Two-photon fluorescence imaging was performed with a femtosecond pulsed Ti:Sapphire laser (Cameleon Ultra II, Coherent) tuned to 840 nm coupled into an Ultima laser scanning head (Ultima scanning head, Bruker), mounted on an Olympus BX61WI microscope, and equipped with a water-immersion objective (60X, 1.1 numerical aperture, Olympus Optical, Tokyo, Japan). Cell morphology was visualized using fluorescence imaging of patch-loaded Alexa 594 (10 μM). Single action potential-induced Ca^2+^ transients (sAPCaTs) were recorded using the calcium indicators OGB-5N, Fluo-5F and Fluo-4, using either rapid line scan imaging (∼10 μm at 0.76 ms per line) or parked beam detection, also referred to as spot detection (Nakamura et al. 2015). Linescan imaging of presynaptic boutons was performed by bisecting the short axis of the bouton. Total laser illumination per single sweep lasted 200 ms. Fluorescence recordings started at 15 min for GC and 30 min for SC (Figure 1 and Figure S1) or after >20 min for GC and >60 min for SC (Figure 2G, Figure3, Figure 4 and figure 6) after establishing the whole-cell configuration. On average, 10 sAPCaTs were recorded per bouton at 0.33 Hz, and a maximum of 3-4 boutons was recorded per cell. The two-photon point spread function was estimated by imaging 100 nm fluorescence beads (PSF_XY_ = 328 ± 11; PSF_z_ = 1132 ± 8, 810nm). The PSF was enlarged by underfilling the objective back pupil with a collimated laser beam (PSF_XY_ = 0.82 μm ± 0.1; PSF_z_ = 10.2 ± 0.3 μm). Fluorescence light was separated from the excitation path through a long pass dichroic (660dcxr; Chroma, USA), split into green and red channels with a second long pass dichroic (575dcxr; Chroma, USA), and cleaned up with band pass filters (hq525/70 and hq607/45; Chroma, USA). Fluorescence was detected using both proximal epifluorescence and substage photomultiplier tubes: multi-alkali (R3896, Hamamatsu, Japan) and gallium arsenide phosphide (H7422PA-40 SEL, Hamamatsu) for the red and green channels, respectively. Two-photon spot-detected fluorescence signals were filtered at 10 kHz using an 8-pole Bessel filter (Frequency Devices), digitized at 100 kHz, then filtered offline (see below).

### Analysis of Ca^2+^ transients

sAPCaTs were constructed from line scan images by averaging the fluorescence over visually identified pixels corresponding to width of the bouton, resulting in a single fluorescence trace as a function of time. The background fluorescence (F_back_) was estimated from the average pixel intensity of those pixels not on a labeled structure, and subtracted from the fluorescence trace. For spot detection, average calcium transients (30-60 sweeps) were corrected for background florescence estimated by the average fluorescence obtained while placing illumination spot in a location away (approximately 30 μm) from the visualized bouton. The background corrected trace was then converted to ΔF/F(t) according to equation 1:.

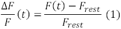

where F_rest_ is the time averaged (20 ms window) fluorescence before stimulation and F(t) is the time-dependent fluorescence transient. Traces were further analyzed if amplitude of ΔF/F trace was 3× larger than SD of baseline. To estimate the amplitude, rise and half-decay time of APCaTs, we fit averaged ΔF/F traces with the following equation (Nielsen et al., 2004), a least-square algorithm implemented in IgorPro (Wavemetrics):

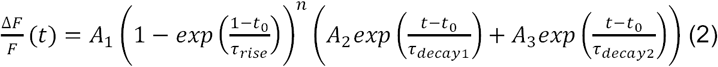

For estimating duration of calcium entry, two-photon spot detected OGB-5N (500 μM) sAPCaTs from each bouton was peak normalized and aligned to the 20% rise time using equation 2. The duration of the Ca^2+^ entry was estimated from a Guassian fit to the first derivative of the sAPCaT. To maximize the signal-to-noise ratio when analyzing population average traces we determined the maximum offline filtering that would not alter estimated duration of calcium entry. For that we applied different values of a binomial smoothing function corresponding to different filtering frequencies (1-10kHz) to population average traces and estimated duration of calcium entry. 10-90 rise times showed little variation for higher bandwidths, and progressively increased with increased filtering. The final values (5 khz for GC and a 3khz for SC) used represent corner frequency in which SNR was improved, but little effect on the rise time was observed (<10%).

### Optical Fluctuation analysis

Estimation of number and P_open_ of VGCCs present in SC and GC boutons was made using fluctuation analysis performed as described previously (Sabatini and Svoboda, 2000). In brief, if VGCC gating is governed by binomial statistics, then the coefficient of variation of the shot-noise subtracted fluorescence variance, is given by:

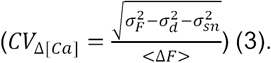

Where 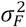 is the total variance, 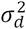 is the dark noise and 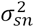 represents the shot noise. The average fluorescence transient and its variance were calculated from the unnormalized fluorescence from 60-120 trials (delivered at 0.33 Hz). In SCs sAPCaT recordings were performed >30 min after establishing the whole-cell configuration. In GCs, we used a fast loading approach in which they were briefly patched for 60-120 sec with an intracellular solution containing Alexa-594 (10 μM) and Fluo-5F (400 μM) after which pipette was carefully removed. The recording was discontinued in cells where somatic calcium levels increased (approximately 3X) after pipette removal. Recordings were initiated 15 min later to allow dye equilibration.

For SC and GCs, dark noise was measured with shutter closed and shot noise was estimated by imaging closed-end pipettes (made using a Microforge-Mf-900 Narishige) filled with the Fluo-5F, positioned at slice surface and varying laser intensity. Variance was calculated for 10 ms baseline period and for 10 ms after the peak in calcium transient. If baseline variance was more than 1 SEM of the expected shot noise variance, bouton was removed from analysis. We estimated coefficient of variance in control conditions and in the presence of baclofen (10 μM), a GABA_B_ receptor agonist that is known to reduce P_open_ of VGCCs (Sabatini and Svoboda, 2000). P_open_ and number of VGCCs (N) were calculated using the following equations:

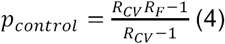

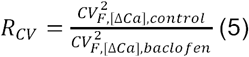

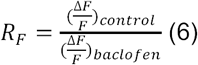

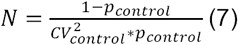

Error in *p*control calculation was estimated using a general error propagation equation, where for any function f(x, y), the uncertainty of the calculation (σ _f_) is given by:

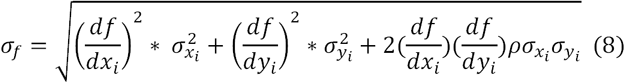

in which the last product accounts for the correlations between, X and Y. In the case of uncertainty propagation for R_CV_ quantification (equation 5), the correlation product refers to that between 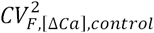 and 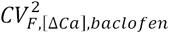. For single boutons both CV^2^ values should be correlated since baclofen has mainly an effect on *p*_open_, and not on N (Sabatini and Svoboda, 2000), but we were unable to record from a single bouton in control and baclofen. However, if we compare the slope of the plot 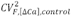 as a function of (ΔF/F)^-1^_control_, to that of 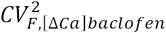 as a function of (ΔF/F)^-1^, we observed a similar correlation coefficient (0.82 for GC and 0.75 for SC). Replacing the product N·p_open_ by ΔF/F in equation 9 suggests that the observed correlations between 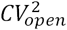 and (ΔF/F)^-1^ indicate similar p_open_ values across different boutons.

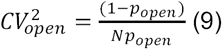

Since N appears to be the main contributor to the measured 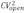 we used the estimated correlation coefficient between 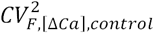 and (ΔF/F)^-1^ as an approximate value for the true correlation between 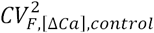 and 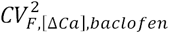 in equation 8.

### Confocal imaging

To determine bouton dimensions, SC and GC were patch loaded with Alexa-594 (40 μM) and imaged using confocal laser scanning optics (Ultima scan head, Bruker). Alexa-594 was excited using a 590 nm laser collimated and adjusted to overfill a 1.1 NA 60X objective (LUMFLN60XW, Olympus). Emitted fluorescence was descaned, and aligned through a 60 μm pinhole, placed on a conjugate image plane (corresponding to ∼0.5 Airy units). The resulting point spread function size was estimated from the FWHM of line profiles along images of 100 nm fluorescence beads (xy = 212 ± 7 nm, z = 672 ± 32 nm). Fluorescence emission from Alexa 594 was filtered with a 605LP filter (Chroma) and detected with a multi-alkali PMT (3896, Hamamatsu Photonics). To determine the bouton volume we first estimated its width and length using the FWHM of fine profiles of the fluorescence intensity of the bouton along the major and minor bouton axes. The bouton volume was the calculated using the equation for an ellipsoid: V_bouton_ = 4/3 * π * R_major_ * R^2^_minor_.

### EGTA-AM calibration experiments

To generate EGTA calibration curves, SC and GC were whole-cell patch-clamped with intracellular solution containing 200 μM OGB-5N together with different amounts of EGTA (0,1, 2, 5 and 10 mM). Free Ca^2+^ was maintained constant (50 nM) though addition of CaCl_2_calculated using Maxchelator). After an equilibration period (>20 min GC, >60 min SC), 30 sAPCaTs were recorded using two-photon detection with enlarged PSF.

To estimate the amount of EGTA loaded in the boutons the decay time of sAPCaTs recorded in boutons after fast loading of 200 μM OGB-5N (see above) and 20 minute exposure to 100 μM EGTA-AM and compared to the calibration curve generated from whole cell dialysis with different EGTA concentrations. To evoke action potentials a second pipette was placed in close proximity of the visually identified axon. We used a transmitted light PMT mounted after the Dodt tube to acquire a laser-illuminated contrast image simultaneously with the 2PLSM image. This dual imaging mode was used to position stimulation electrodes close to the axon. In GC, stimulation intensity was settled while imaging OGB-5N fluorescence in single boutons. Intensity (4-40 V, 50 μsec) was progressively increased until reliable APCaTs were induced. Because of the small peak amplitude of sAPCaTs recorded from SCs, we adjusted stimulation intensity by monitoring AP generation in SC soma using cell-attached recordings. EGTA-AM was prepared in DMSO and diluted in ASCF every day. Stock solutions were used for a maximum of 2 weeks to minimize degradation.

In comparison to the fast loading approach, whole-cell dialysis itself did not alter the decay of GC sAPCaTs (14.4 ± 1.3 ms, n = 21 in whole-cell, 14.7 ± 2.1 ms, n = 15, using fast loading; p=0.90, Mann-Whitney test; Figure 4I). However, for SC boutons, the obtained half-decay under fast loading conditions was significantly smaller (3.4 ± 0.5 ms, n=15, p < 0.0001, Mann Whitney test) than the control half-decay (0.1 mM EGTA value) obtained in whole-cell configuration, consistent with the wash-out of slow binding endogenous calcium buffers (e.g. parvalbumin, (Eggermann and Jonas, 2011).

### Monte Carlo simulations of Ca^2+^ reaction-diffusion and vesicle release

Stochastic simulations of Ca^2+^ reaction-diffusion dynamics and vesicle release probability were performed using MCell 3.4 (http://mcell.org; Kerr R. et al., 2008). The overall model comprised Ca^2+^, VGCCs, vesicular sensor, as well a fixed and mobile (EGTA, ATP, calretinin and parvalbumin) buffers. The size of the simulated compartment was approximated by an elliptic cylinder based on boutons dimensions estimated from confocal imaging of SC and GCs (Figure S1; Table S1). The walls of the compartment were set as reflective for all diffusants. The VGCCs were located on one of the flat surfaces. Labelling efficiency was estimated by dividing the obtained total number (median) of gold particles per bouton with the median number of VGCCs estimated by optical fluctuation analysis (19% for GC and 46% for SC). For SC simulations, the PRM model was used. VGCCs we placed in a cluster with a mean channel NND of 15 nm, corresponding to 17.6+-/0.5 nm for 16 VGCCs (mean +/-SEM) after accounting for a labelling efficiency (estimated from null model simulations), and was close to the experimentally obtained NND of 20.4 ± 7.0 nm (mean ± sd; Figure S3J).

AP-mediated Ca^2+^ entry was simulated using stochastic channel gating scheme (see below). Reaction-diffusion simulations included 4 mM of a low affinity (K_d_ = 100 μM) fixed buffer (Nakamura et al., 2015), giving a buffer capacity of 40, similar to that estimated in Figure 1. ATP was added at a concentration of 200 μM, corresponding to the estimated free concentration obtained from a total ATP concentration of 1.5 mM (Rangaraju et al., 2014) in the presence of 700 μm free Mg^2+^ and 50 nm resting [Ca^2+^], as calculated using Maxchelator. The concentration and rate constants of the endogenous mobile buffers, parvalbumin (500 μM) in SC and calretinin (160 μM) in GC were defined in agreement with previously published values (Eggermann and Jonas, 2011; Schmidt et al., 2013), and listed in Table S1. To calculate EGTA inhibition in P_v_ a separate set of simulations was run in the presence of EGTA for both GC and SC. For each simulation condition, the P_v_ was computed as average number of release events over 2000-5000 independent trials. To reduce computational time simulations were performed using the 2ms window which is sufficient to account for 80% of total release events for the slower EZM (50 nm), tested against a 4 ms simulation (Figure S5). Deconvolution of GC-SC EPSCs with quantal responses (Abrahamsson et al., 2012) revealed that the peak of evoked EPSC, which was used to estimate release probability (Baur et al., 2015) at GC synapses, occurs at a similar percentage (82%) of the release integral. The simulator was run on the TARS cluster of Institut Pasteur, Paris, France.

### Simulation of VGCC gating

Individual VGCCs were modeled using Hodgkin & Huxley four-state gating model (Borst and Sakmann, 1998):

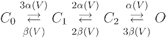

where *C*_}_ denotes a closed state, *C*_;_ and *C*_J_ are two intermediate closed states and *O* is an open state. The voltage-dependent rate constants *α*(*V*) and *β*(*V*) are calculated according to the following formulas:

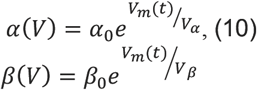

where *V*_*m*_*(t)* is the membrane potential generated by a modified AP-waveform measured at the calyx of Held (Nakamura et al., 2015) and the resting membrane potential was set to −75 mV. The small hyperpolarization phase was set to the resting membrane potential. *α*_*0*_, *β*_*0*_, *V*_*α*_, *V*_*β*_ are gating parameters that were chosen such that resulting peak of single channel P_open_ (0.285) and half duration (250 μs) of a single channel calcium current match with experimentally observed values (Figure 2 and Figure 4). The single channel current was calculated: *I*(*t*) =^*g*^/_2*e*_ (*V*_*m*_(*t*) - *E*_*rev*_), where *g* is the single channel conductance, *e* is the elementary charge and *E*_*rev*_ = 45 mV is the reversal potential. A five-state release model was used to simulate P_v_ (Wang et al., 2008). The remaining parameters are listed in Table S1.

The broadened AP (Figure 6) was constructed as follows: the hyperpolarization component of the control AP waveform was set to the resting membrane potential then modified waveform was fitted with a double Gaussian function using equation 11:

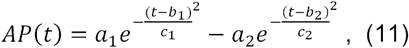

and parameters *a*_*1*_=73.48, *b*_*1*_=0.4476, *a*_*2*_=52.31, *b*_*2*_=0.5529. To generate an *I(t)* that was increased in integral by 50 and 100% we used the following parameters in equation 11: c1=0.0955 and c2=0.1329, and c1=0.0735 and c2=0.1022, respectively. In order to keep depolarization of the AP intact the rising phase of the broadened curves calculated by (11) were replaced with the depolarization phase of initial AP. Final waveforms are shown in Figure 6.

### Statistical analyses

Data are presented as average ± SEM unless otherwise indicated. Statistical tests were performed using a non-parametric Wilcoxon-Mann-Whitney two-sample rank test for unpaired, a Wilcoxon signed-rank test for paired comparisons, and a Kruskal-Wallis test followed by Dunn’s post hoc test for multiple comparisons. Linear correlations were determined using a Pearson test. For particle-based reaction-diffusion simulations (Figure 6), data is presented as mean ± SEM and a Chi-squared test was used to detect statistical difference between the simulated release probabilities. In the case of comparing P_v_ between two conditions (X and Y) the error (σ) was propagated according to the following formula:

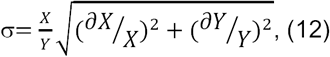

### SDS-digested freeze-fracture replica-labeling (SDS-FRL)

Five male C57Bl6j mice (P35–45) were deeply anaesthetized and transcardially perfused with a fixative containing 2% PFA and 0.2% picric acid in 0.1 M PB for 15 minutes. 80 μm thick sagittal sections from the cerebellar vermis were cut with a vibratome, cryoprotected in 30% glycerol. Tissue samples from WT and cKO mice lacking the Cav2.1 subunit from GCs were kindly provided by Ryuichi Shigemoto (IST Austria) and Melanie D. Mark (Department of Zoology and Neurobiology, Ruhr-University Bochum). Replicas were prepared as described previously (Kirizs et al., 2014). The replicas were washed and blocked with 0.5 or 5% BSA in Tris buffered saline (TBS) for 1 hour followed by an incubation in a solution of the following primary antibodies: rabbit anti-Cav2.1 (1:500; Synaptic Systems, Cat No. 152 203, RRID:AB_2619841; the specificity was tested in cerebellar tissue samples from Cav2.1 cKO mice, Figure S2), guinea pig anti-Cav2.1 (1:50, Frontier Institute, Cat No. VDCCa1A-GP - Af810, RRID: AB_2571851; the specificity of which was demonstrated in (Holderith et al., 2012), rabbit anti-Cav2.2 (1:400; Synaptic Systems; Cat No.: 152 303; RRID: AB_2619844; specificity of the reaction with this antibody was verified in (Lenkey et al., 2015), guinea pig anti-VGAT (1:1500; Synaptic Systems, Cat. No. 131 004, RRID: AB_887873), rabbit anti-Munc13-1 (1:400, Synaptic Systems, Cat. No. 126 103, RRID:AB_887733). Replicas then were washed and incubated in a solution containing the following secondary antibodies: goat anti-rabbit IgGs coupled to 5 or 10 nm gold particles (1:75 or 1:100; British Biocell), goat anti-guinea pig IgGs coupled to 15 nm gold particles (1:100; British Biocell) or donkey anti-guinea pig IgGs coupled to 12 nm gold particles (1:25, Jackson ImmunoResearch). Replicas were rinsed in TBS and distilled water before they were picked up on copper parallel bar grids and examined with a Jeol1011 EM. To determine the Cav2.1 subunit content of boutons, electron micrographs of VGAT-negative (putative GC) and VGAT-positive (putative SC) boutons containing Cav2.1 labeling were randomly acquired in the molecular layer. VGAT-negative boutons with more than two putative AZs were excluded from the analyses (putative climbing fiber boutons). The AZs of VGAT-negative boutons were delineated based on the underlying high density of intramembrane particles. Nonspecific labeling was determined on surrounding exoplasmic-face plasma membranes and was found to be 0.8 and 1.5 gold particle / μm^2^ for the rabbit and the guinea pig anti-Cav2.1 antibodies, respectively, and 0.8 and 1.1 gold particle / μm^2^ for the anti-Cav2.2 and anti-Munc13.1 antibodies, respectively.

### Gold particle analysis

Coordinates of the immunogold particles and corresponding AZ perimeters were extracted from EM image. Spatial organization of immunogold particles in presynaptic AZs was analyzed on the population of AZs for each synapse type using mean nearest neighbor distance (NND) and a Ripley analysis (Besag, 1977; Ripley, 1977), while single AZ analysis was performed using Ripley’s functions.

Mean NND between gold particles at each AZ is a local density measure that can be used to detect potential deviations from a null model within the AZ area, i.e. a homogeneous distribution of random particle locations.

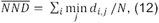

where *N* - number of gold particles within the AZ, and *d*_*i,j*_ is the Euclidian distance between points *i* and *j*. Mean NND distributions were compared to null model NND distributions generated from 1000 Monte Carlo simulations of random particle placement within the AZ, using Mann-Whitney test with Bonferroni correction.

Ripley K-function is used to examine whether particle distributions are clustered or dispersed over a range of spatial scales. Ripley’s function reports the normalized density at different spatial scales. The Ripley’s function with edge correction is defined as:

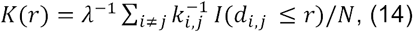

where *λ* is density of a point pattern; *k(i,j)* is a fraction of a circumference of radius *r* with center at point *i*, passing through point *j* and that lies within the boundary (edge correction term); function *I* is an indicator function (takes value of 1 if its operand is true and 0 otherwise); *N* is number of points in the pattern. We finally used a variance stabilized and boundary corrected version of the Ripley’s K function and r is the radius; Diggle 2003, see also (Kiskowski et al., 2009), which we call the H-function:

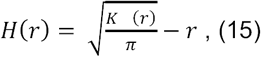

H-function analysis was performed for each gold particle pattern in the population (SC or GC PPs) and corresponding patterns (n=5000) generated using the null hypothesis. For each population the *H(r)* functions were pooled according to:

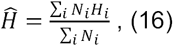

where *N*_*i*_ - number of points in the *i*th sample and the *H*_*i*_ – H function of the *i*th sample (Diggle,2003). The 99% confidence envelops (CE) for pooled *H(r)* function were calculated based on the H functions calculated for the whole set of generated patterns. Departure from the null model was tested using Maximum Absolute Deviation (MAD) test (see below); for each population (GC or SC), based on pooled quantities.

<u>Single sample</u> Ripley H-function analysis was computed for each AZ pattern within the GC or SC population. The 99% confidence envelops (CE) for H-function were calculated based on 5000 patterns generated using Monte Carlo simulations of the null hypothesis. MAD test was applied to detect departure from the null model.

### Maximum Absolute Difference (MAD) test

Statistical departure of the Ripley analysis from the null model was determined by the (MAD test, (Baddeley, 2014). The MAD test is an inferential statistical tool that quantifies the departure of the summary function (nearest neighbor distribution, spatial correlation function or Ripley’s K or H functions) from the null model, and is based on the estimation of the maximum deviation between the functions. The maximum deviation is computed as follows:

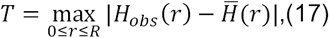

where *R* is an upper limit of the interaction distance (distance to AZ perimeter), 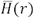 is the expected summary function for the null model and *H*_*obs*_(*r*)for the observed pattern. 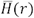 was estimated based on m (5000) simulation of the null model as follows:

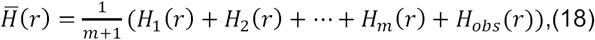

where *H*_*i*_(*r*) is the H-function computed for the *i*th pattern generated under the null model. Deviation between the summary function of observed pattern and each simulated pattern under the null model was calculated as: 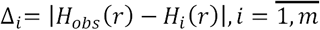. Then the null hypothesis was rejected if for the predefined confidence (99%, a common criteria; Tian et al., 2007) level ^*l*^/_*m*+ 1_ if the following holds: *T* > *Δ*_*l*_. The p value was computed as

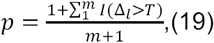

### Bivariate Ripley’s function

The Bivariate Ripley’s function allowed us to study relative distribution of two different particle types, representing Cav2.1 and Munc13-1. Assuming two types of particles (*i* and *j*) in a given pattern, then the bivariate Ripley K-function is (Hanisch, 1979):

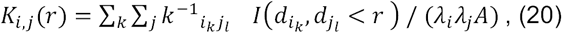

+where *λ*_i_ is density of points type *i* (*λ*_j_ of points type *j*); *A* is area of the window; *k(i,j)* an edge correction factor, analogous to the univariate faction case; function *I* is an indicator function; *d* is a Euclidean distance between points in the pattern of the same type. The variance normalized bivariate Ripley’s function is function is:

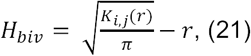

The 99% CE for *H*_*biv*_ function were calculated from 5000 patterns produced using Monte Carlo simulations of the null model, generated for this analysis by merely reassigning, randomly, each original particle as either as a Munc13-1 or Cav2.1 particle. In order to pool different *H*_*biv*_ functions from the different patterns across independent AZs, we used the 99% CE estimated from the Monte Carlo simulations to scale the individual *H*_*biv*_ function for each AZ, such that the 99% CI appears as +/-1 on graph.

### Null model simulations

To generate spatially random patterns of particles within AZs, particle locations were generated by uniform-random sampling of the Cartesian coordinate space within a the minimum sized rectangle that fully encompasses the AZ. Particle locations generated outside of the AZ were discarded. Points that were closer than a chosen distance were excluded to respect the obtained minimum distance between the labeled Cav2.1 in EM data which was, 5.3 nm in GC and 7.1 nm in SC. The total number of generated points was defined by either the number of particles observed in the EM sample or 34 points in case of the “idealized” GC analysis (Figure S3A-C).

### Exclusion Zone point pattern simulation

SVs were randomly placed within the AZ area, determined from EM images described above, in a manner similar to simulated particle patterns. Locations that were closer than 20 nm corresponding to SV diameter were discarded. The number of SVs used for each active zone (N_SV_) was calculated from 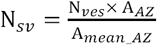, where A_*AZ*_ is the area of a given AZ, A_*mean*_*AZ*_ = 0.09 *um*^2^and is the mean GC AZ area, and N_*ves*_ = 8 +/-2, the average number of vesicles per AZ based (Xu-Friedman et al., 2001). To simulate Cav2.1 particle patterns, points (the same number of measured gold particles for a given AZ) were randomly distributed within the measured area of the same AZ, as for the null model, except particles falling at a distance less than an exclusion radius (r_ez_) were discarded.

### Cluster point pattern simulation

Point coordinates representing cluster centers were first randomly placed in a circular AZ (area=0.09 μm^2^ (Figure S3), as in the null model. N_"_ cluster center points that were closer than a minimum distance (d_*cl*_) were considered for further analysis. Point clusters were generated around each *N*_*c*_ by sampling their coordinates from a Gaussian distribution where the mean is the cluster’s center and the SD set such that the distance between the elements of the single cluster is not closer than d_*p*_*min*__ and not further than d_*p*_*max*__. The number of points per cluster, N_p_ points was set such that N_c_ ∗ N_p_ = 180 simulated particles –similar to the total number of VGCCs estimated to be present in GC boutons by optical fluctuation analysis (Figure 2). Parameters are listed in Table S2.

### Bivariate NND null model simulation

For each AZ we randomly generated a number of particle locations that corresponds to the number of Munc13-1 particles, but were not closer than 10.2 nm. Similarly, we randomly generated particle locations corresponding to Cav2.1 that were not closer than 7 nm, and not closer than 8.3 nm to Munc13-1 particles. The minimum distance between the particles are estimated from the data.

### Bivariate EZ model simulation

The EZ point pattern corresponding to Cav2.1 labelling was generated as described above using minimum distance between Cav2.1 points of 7 nm. Subsequently a number of points representing Munc13.1 were randomly placed in the measured AZ area. Then were rejected such that simulated Munc13-1 particles formed one of three patterns: 1) particles remained randomly placed inside the EZ, 2) particles remained outside of the EZ, or 3) Munc13.1 were randomly distributed throughout the AZ. Munc13-1 points could not be closer than a minimum distance to other Munc13-1 points (10.2 nm) and to Cav2.1 points (8.3 nm) estimated from EM data (Figure 7). The final number of points generated for each Munc13-1 pattern corresponds to the total number of Munc13-1 particles observed for each AZ.

All calculations were performed on the desktop computer, process: Intel Xeon CPU E5-1620v3 3.50 GHz, 16 GB RAM, 64 bit operation system Windows 7 Ultimate, Matlab R2017b.

### Clustering of gold particles with DBSCAN

To determine the number of gold particle clusters of Cav2.1 and Munc13-1 gold particles SC boutons DBSCAN (Ester et al., 1996) was used, which is a density-based clustering algorithm. We used either Matlab custom code or a Python-based open source software with a graphical user interface, GoldExt (Szoboszlay et al., 2017). The DBSCAN requires two user-defined parameters: ε (nm), which is the maximum distance between two localization points to be assigned to the same cluster, and the minimum number of localization points within a single cluster. The epsilon parameter was determined from a set of DBSCAN analyses for different epsilon values (with a minimum number of points per cluster of one). We selected 50 nm as it approximates the distance at which the deviation is maximum between the data and randomly generated patterns (46 nm; Figure S3D). Since the density of single gold particles per SC bouton (∼ 2.5/μm^2^) is on the order of the background labelling density (1/μm^2^) we used a minimum number of 2 particles to define a cluster for subsequent analyses. This rejected 9% of total particles from the analysis.

---

**Figure S1:**
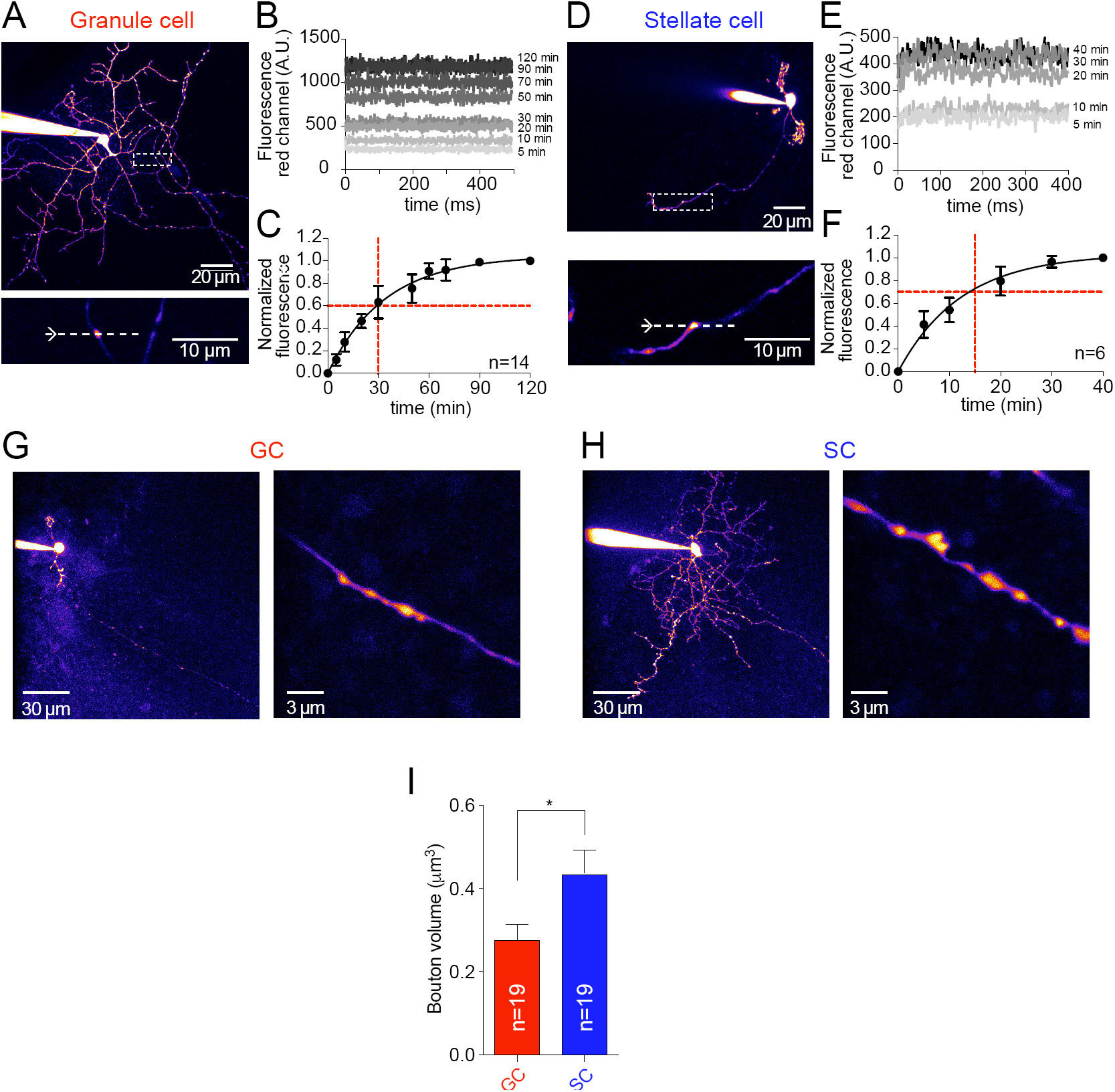
Quantification of loading times and volumes of GC and SC boutons. **(A)** *top*: 2PLSM MPI image of patch-labelled GC (Alexa-594-10 μM). *bottom*: Axonal region corresponding to dashed box from upper image with imaged bouton. **(B)** Time series traces are mean (10 sweeps) fluorescence of individual Alexa-594 images over 0.7 μm obtained from the bouton illustrated in A at different time points after entering whole-cell. **(C)** Population average (n=14 cells) of normalized (to value obtained at 120 min) average Alexa-594 fluorescence for different time points obtained at different time points after entering whole-cell. Black line represents a single exponential fit. Dashed red lines indicates the time used in whole cell configuration to image calcium transients. Error bars represent ± SEM **(D-F)** Same as A-C but for SC. **(G)** Confocal image (MPI) of a patch-loaded GC (Alexa-594-10 μM; top) and of magnified view of the axon. **(H)** Confocal image (MPI) of a patch-loaded SC (Alexa-594-10 μM; top) and of magnified view of the axon. **(I)** Summary plots of estimated bouton volumes for GCs and SCs. Error bars represent ± SEM. (p<0.05, Mann-Whitney test).

**Figure S2:**
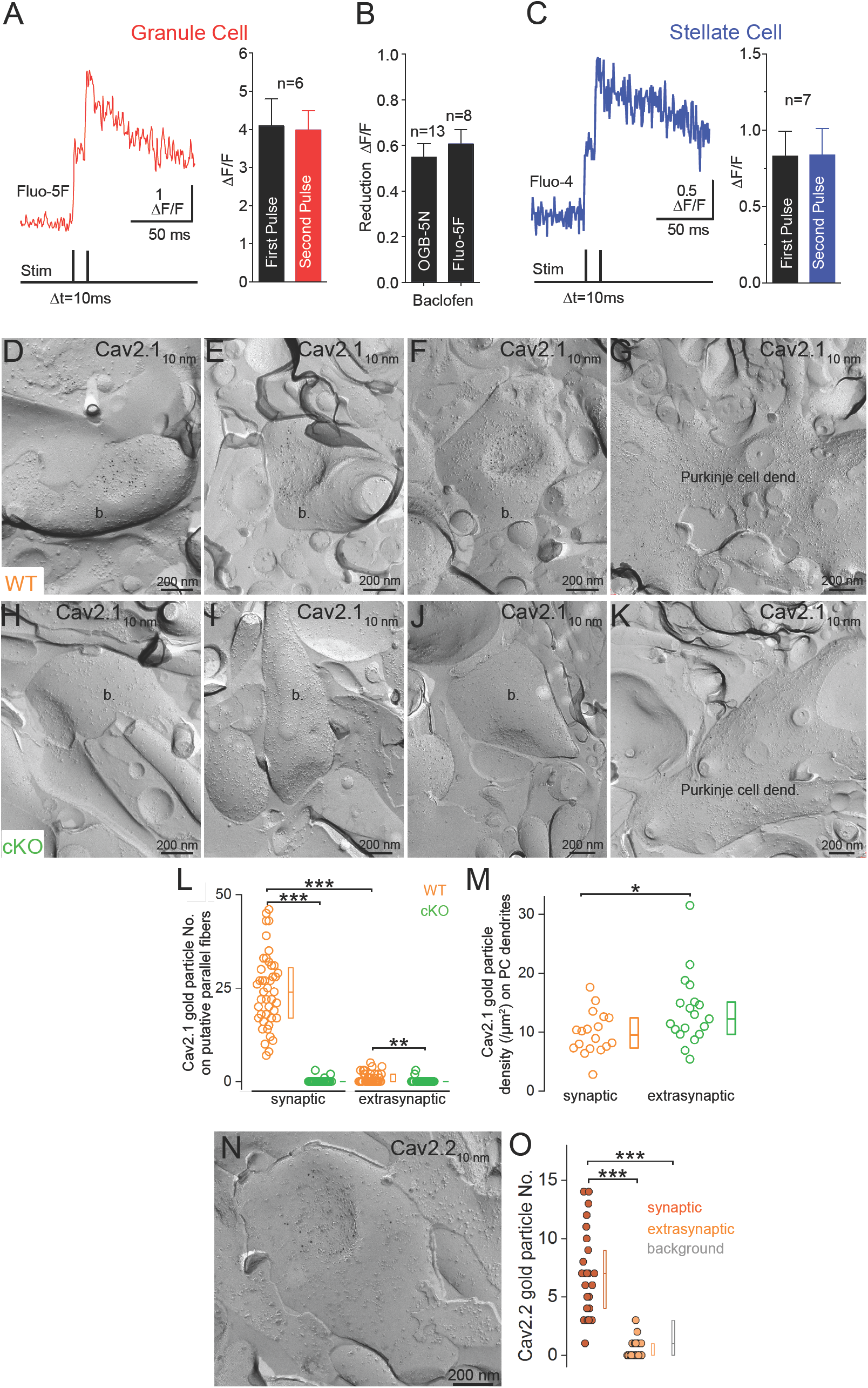
Linear behavior of Fluo-5F and Fluo-4 under working conditions used for optical fluctuation analysis and specificity of immunoreactions performed with the rabbit anti-Cav2.1 antibody. **(A)** *Left*: Averaged line scan image (from 5 images) of GC sAPCaTs (Fluo-5F) recorded in the fast loading configuration in response to a paired-pulse extracellular stimulation of PFs. *Right:* Peak amplitude (n=6 boutons) of sAPCaTs for the first and second pulse of the paired stimulation. Box indicates mean ± SEM. **(B)** Summary plot of inhibitory effect of 10 μM Baclofen in peak amplitude of sAPCaTs recorded with either OGB-5N (100 μM, n=13) or Fluo-5F (fast loading, n=8). Box indicates mean ± SEM. **(C)** *Left*: SC APCaTs (Fluo-4, 200 μM) recorded in response to a paired-pulse stimulation evoked by intracellular current injection. *Right:* Peak amplitude (n=7 boutons) of sAPCaTs for the first and second pulse of the paired stimulation. Box indicates mean ± SEM. **(D-F)** In putative GC boutons (b.) of a WT mouse gold particles labeling the Cav2.1 subunit are enriched in AZs indicated by the high density of intramembrane particles. **(G)** In the same mouse, a Purkinje cell dendrite (dend.) contains scattered gold particles labeling the Cav2.1 subunit. **(H-K)** In the cKO mouse, gold particles labeling the Cav2.1 subunit are absent from putative GC boutons (J-L), while Purkinje cell dendrites were immunopositive (K). **(L)** The number of gold particles labeling the Cav2.1 subunit in AZs and extrasynaptic membrane segments of GC boutons are significantly higher in WT mouse (orange, n = 44 and 43, respectively) than in cKO mouse (green, n = 28 and 28, respectively; synaptic: p < 0.0001, extrasynaptic: p < 0.003 post hoc Mann-Whitney U tests with Bonferroni correction after Kruskal–Wallis test (p < 0.0001)). Furthermore, in WT mouse there was a significant difference between the Cav2.1 subunit content of synaptic and extrasynaptic compartments (p < 0.0001 post hoc Mann-Whitney U tests with Bonferroni correction after Kruskal–Wallis test (p < 0.0001)). **(M)** The density of Cav2.1 subunit measured on Purkinje cell dendrites is significantly lower in WT (orange, n = 18) than in cKO mouse (green, n = 19; p < 0.05, Mann-Whitney test). Circles indicate individual measurements of boutons (D-F and H-J) or Purkinje cell dendrites (G, K). **(N)** A putative GC bouton is labeled for the Cav2.2 subunit. **(O)** Number of gold particles labeling the Cav2.2 subunit within AZ (syn., n = 25 from 1 mouse), in extrasynaptic membranes (extrasyn., n = 25) and in surrounding EF membranes (background: bg., n = 14). *Post hoc* MW U-test with Bonferroni correction after Kruskal-Wallis test (p < 0.0001) demonstrated a significant difference between the synaptic and background (p < 0.0001), as well as the synaptic and extrasynaptic labeling (p < 0.0001). Circles indicate individual measurements of synaptic or extrasynaptic membranes; the boxes represent interquartile ranges with the horizontal bars showing the medians.

**Figure S3:**
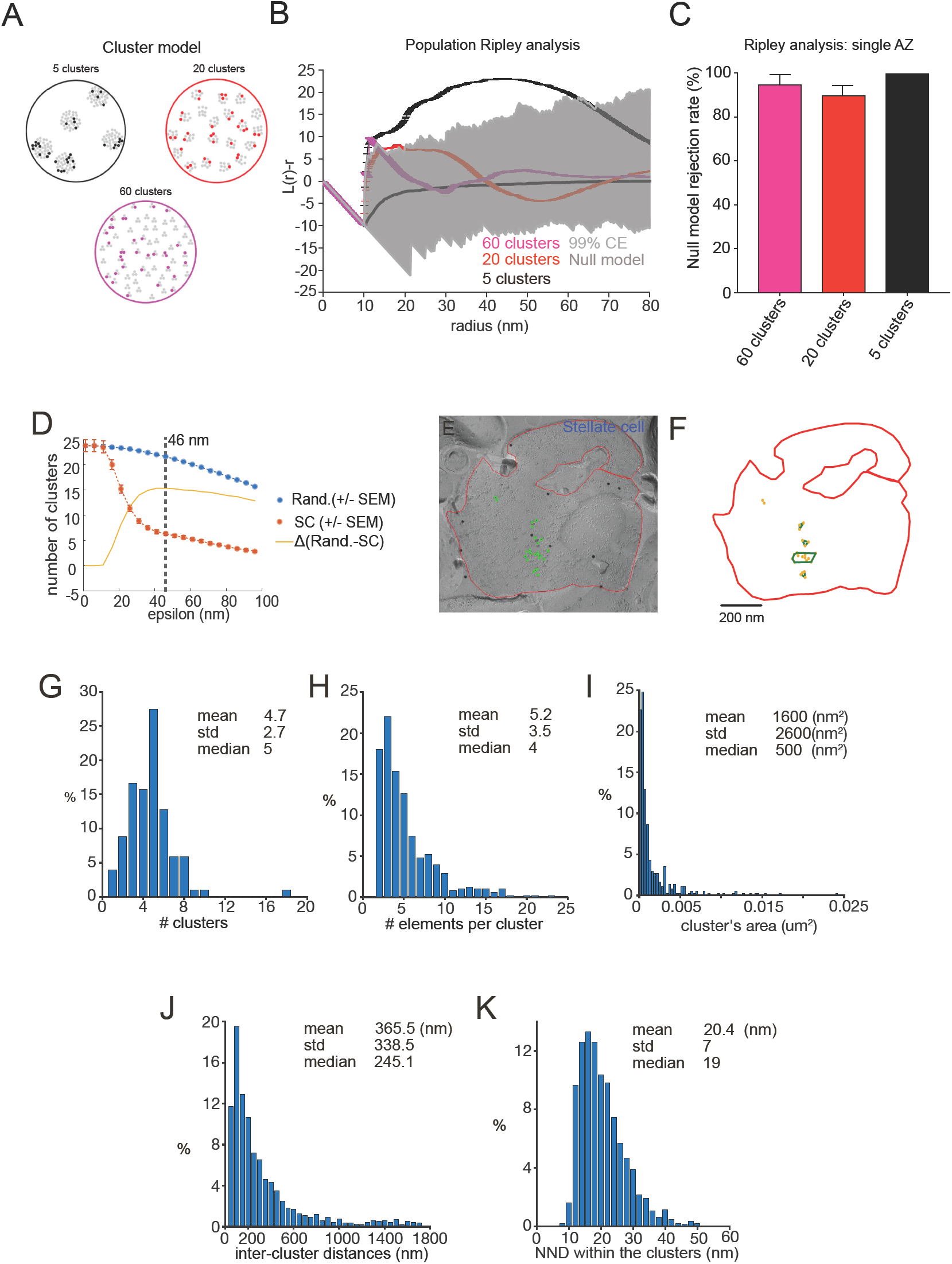
Probing the ability of Ripley analysis to detect clusters of gold particles and cluster analysis in SC boutons. **(A)** Examples of point patterns of three different cluster models that generated either 5, 20 or 60 clusters using 180 points (gray symbols; mean estimated VGCC number/GC AZ). For simplicity, the AZ was defined as a circle with area 0.09 um^2^ (corresponding to mean GC AZ area). 21% of the points were randomly chosen for the analysis (colored symbols), thereby mimicking labeling efficiency of VGCCs by Cav2.1 antibodies conjugated to gold particles. **(B)** Population Ripley analysis of cluster models (5, 20 or 60 clusters). 100 point patterns for each model was generated then the quantity H(r) was computed. The solid gray line is the asymptotical approximation of H(r) function calculated for the null model in the circular AZ. The shaded region indicates the 99% confidence envelope 1000 null model spatial patterns. **(C)** Summary plot showing the percentage of samples in which the H(r) function was not compatible with the null model spatial pattern confidence envelope (CE, from 1000 simulated spatial patterns). Rejection of the null model was determined following a MAD test. The rejection rate was 95 +/- 5% for 40 clusters (pink), 90 +/- 6% for 20 clusters (red) and 100% for 5 clusters (black). Error bars show SEM calculated across simulations for each model. **(D)** Number of clusters estimated in single SC boutons using DBSCAN analysis for different values of epsilon in EM data and in random distributions. Optimal epsilon (46 nm) was defined as the value where the difference in the estimated number of clusters was highest between random and SC data was larger. **(E)** Example of EM image with Cav2.1immunogld labeling in a single SC bouton. **(F)** Cluster identification by DBSCAN analysis of Cav2.1 gold particles from SC bouton (red line) in D. **(G)** Distribution of number of Cav2.1 clusters in the SC boutons (n= 102) in EM data. **(H)** Number of Cav2.1 labeled particles per each identified Cav2.1 cluster in the SC boutons (n= 102). **(I)** Distribution of areas of Cav2.1 clusters in the SC boutons. **(J)** Intercluster distances for Cav2.1 clusters in the SC boutons. **(K)** Distribution of the nearest neighbor distances for labeled Cav2.1 particles within Cav2.1 clusters in the SC boutons.

**Figure S4:**
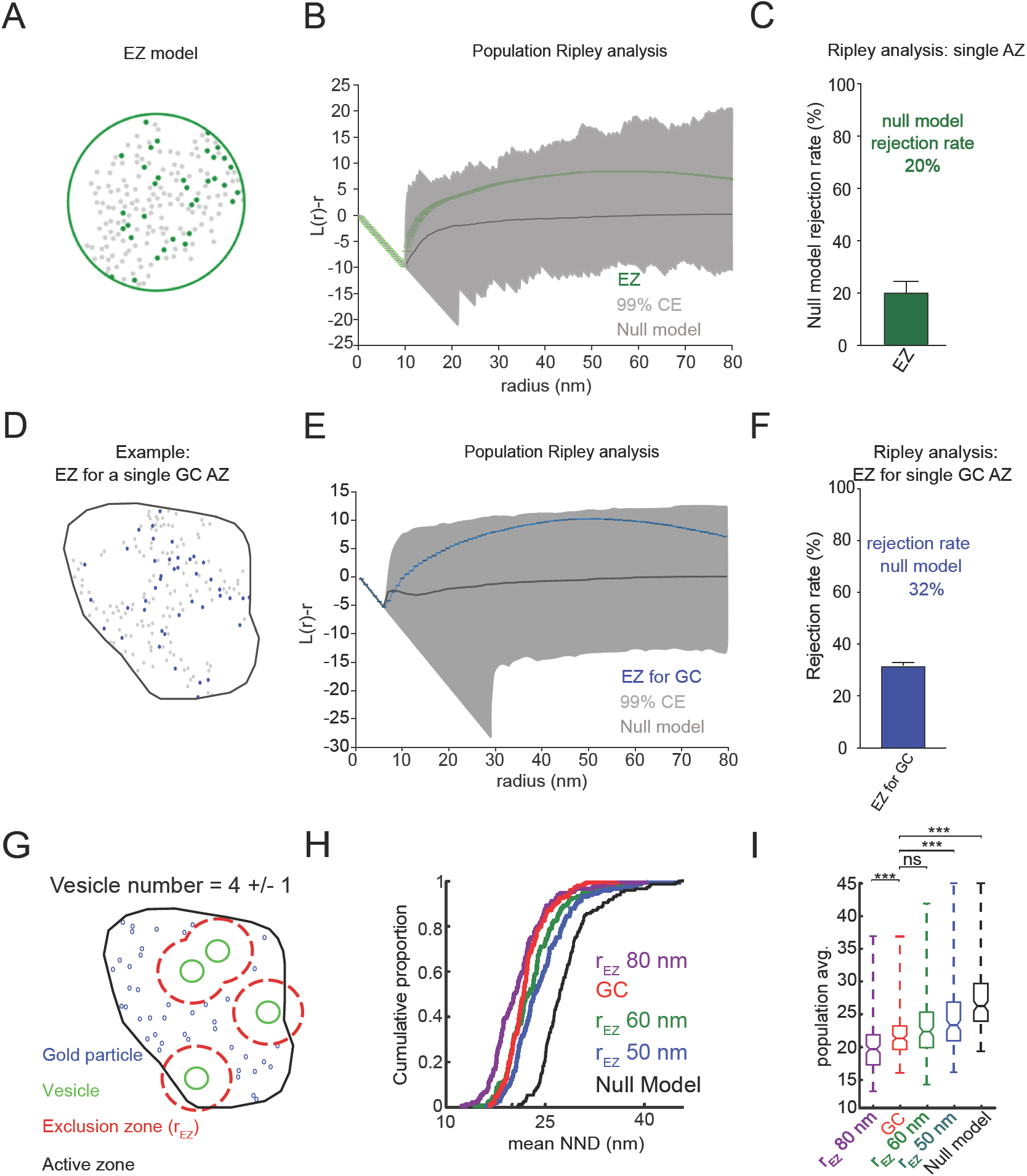
Ability of Ripley analysis to detect modeled exclusion zones (EZs). **A**-Example spatial pattern generated from using the EZ model (EZM; 180 points, grey; 8 +/- 2 docked vesicles). The AZ area was a circle with an area corresponding to the average GZ AZ area (0.09 um^2^). 19% of the overall points (colored points) were randomly chosen for the analysis (mimicking labeling efficiency of gold particles). **(B)** Population Ripley analysis of EZM. The quantity H(r) was computed and pooled for the samples generated under the EZM (n=100) and null model (solid grey line). The shaded region indicates the 99% envelope of values obtained from simulation of the null model for the same area as the AZ. **(C)** Summary plot showing the percentage of samples in which the H(r) function was not compatible with the null model spatial pattern confidence envelope (CE, from 1000 simulated spatial patterns). Rejection of the null model was determined following a MAD test. The number of rejections 20 +/- 5% for the EZM. Error bar represent SEM. **(D)** Example spatial pattern generated for a single GC active zone (AZ) using the EZM (180 points, grey; 8 +/- 2 docked vesicles). 21% of the overall points (colored points) were randomly chosen for the analysis (mimicking labeling efficiency of gold particles). **(E)** Population Ripley analysis of EZM. The quantity H(r) was computed and pooled for the samples generated under the EZM (n=100) and null model (grey line; 1000 simulated spatial patterns were generated per AZ). The shaded region indicates the 99% envelope of values obtained from simulation of the null model for the different GC AZ (n=149). **(F)** Summary plot showing the percentage of samples in which the H(r) function was not compatible with the null model spatial pattern confidence envelope (CE, from 1000 simulated spatial patterns). Rejection of the null model was determined following a MAD test. The number of rejections 32 +/- 2.5 %. Error bar represent SEM. **(G)** Schematic of a single gold particle pattern generated by the EZM for the GC using on average 4 +/- 1 synaptic vesicle per AZ. Docked synaptic vesicles (green circles, r=20 nm) were placed in a uniform random manner within the AZ. An EZ (red dashed lines, r= 50 nm) was drawn around each vesicle and the gold particles placed uniformly in the remaining free space with a minimum distance between two points of 5.3 nm. **(H)** Cumulative distributions of mean NNDs for gold particle distributions in GC AZs (from B), EZM and null models. 1000 simulated patterns were generated for each EZ and hard core model. **(I)** Box plot summary showing EZM with exclusion radii of 60 nm was different from null model mean NND distributions (p << 1e-5; Kruskal-Wallis test followed by Bonferroni post hoc test to account for multiple comparisons). While experimental and EZ (60 nm) mean NND distributions were not (p=0.52).

**Figure S5:**
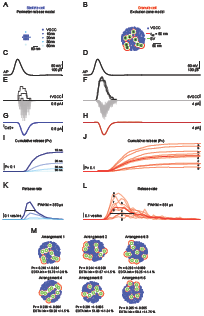
Studying release probability using perimeter release model (PRM; SC) or EZM (GC) with Monte Carlo simulations. **(A)** Model arrangement of VGCC (top) used to probe single vesicle release probability (P_v_) for SCs. The release sensor was placed at four different locations (10, 20, 30 and 50 nm) from the edge of a cluster of VGCCs (16 VGCC-example shown) with an NND of 15 nm. **(B)** EZ arrangement of VGCC and synaptic vesicle used to probe P_v_ in GCs. A total of 8 release sensors, corresponding to 8 synaptic vesicles (green circles, r=20 nm), were randomly placed in AZ with an area of 0.09 um2 corresponding to the average AZ area obtained from EM data. 180 VGCCs were randomly dropped in AZ except in a radius of 50 nm from each release sensor (red dashed lines, r= 50 nm) **(C-D)** Action potential (AP) used to drive gating of modelled VGCCs **(E-F)** Time course of open VGCCs and corresponding calcium current for three different trials. **(G-H)** Mean total calcium current generated by the two different models. **(I)** Cumulative release probability plots obtained for release sensor located at 4 different location from a cluster of VGCCs (Figure 6). **(J)** Cumulative release probability plots for 8 release sensors corresponding to the 8 vesicles displayed in arrangement of Figure 6D. **(K-L)** Vesicular release rate for PRM and EZM. **(M)** Impact of varying arrangement of VGCCs and SV in estimated P_v_ and EGTA inhibition for EZ model (50nm).

**Figure S6:**
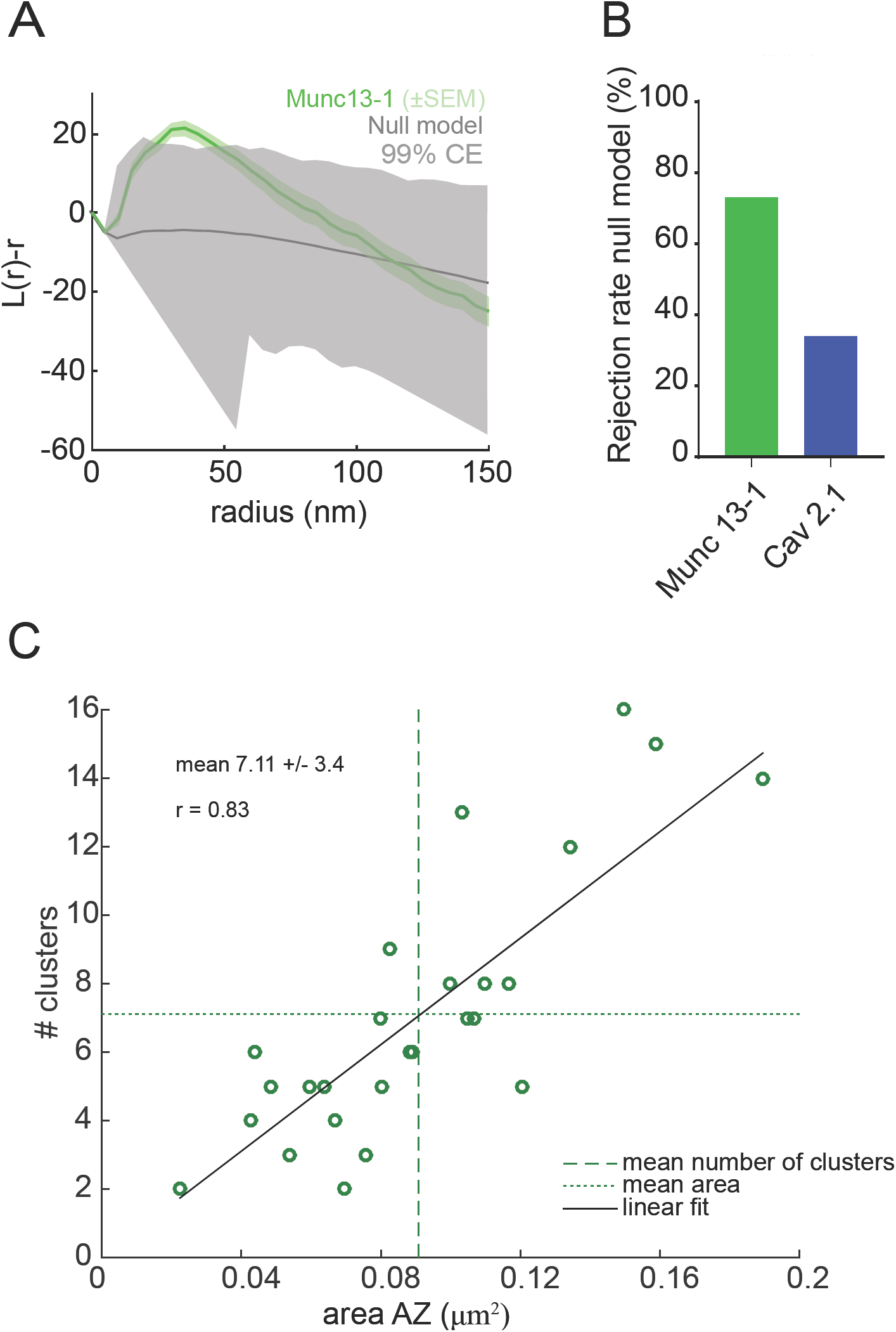
Ripley analysis of Munc13-1gold particle distributions in GC AZ. **(A)** Ripley analysis of gold particle spatial patterns in 26 GC AZs using an edge-corrected H(r) function. Grey lines indicate the 99% confidence envelope obtained from 1000 simulations of the null model for each GC AZ. The quantity H(r) (mean±SEM) as well as the 99 % confidence interval were pooled using a weighted average function. GC AZs are shown with green (n=26). **(B)** Summary plot showing the percentage of samples in which the H_biv_(r) function was not compatible with its paired (same area and number of particles) null model spatial pattern confidence envelope performed using 1000 simulated spatial patterns per AZ area. Rejection of the null model was determined following a MAD test. The null hypothesis was rejected for 71% of Munc13-1 (green) and 35% of Cav2.1 (blue) patterns. **(C)** Number of Munc13-1 gold particle clusters estimated using DBSCAN analysis (epsilon=40 nm and a minimum number of particles per cluster of 1) and AZ area show positive correlation (r, Pearson correlation coefficient).

**Figure S7:**
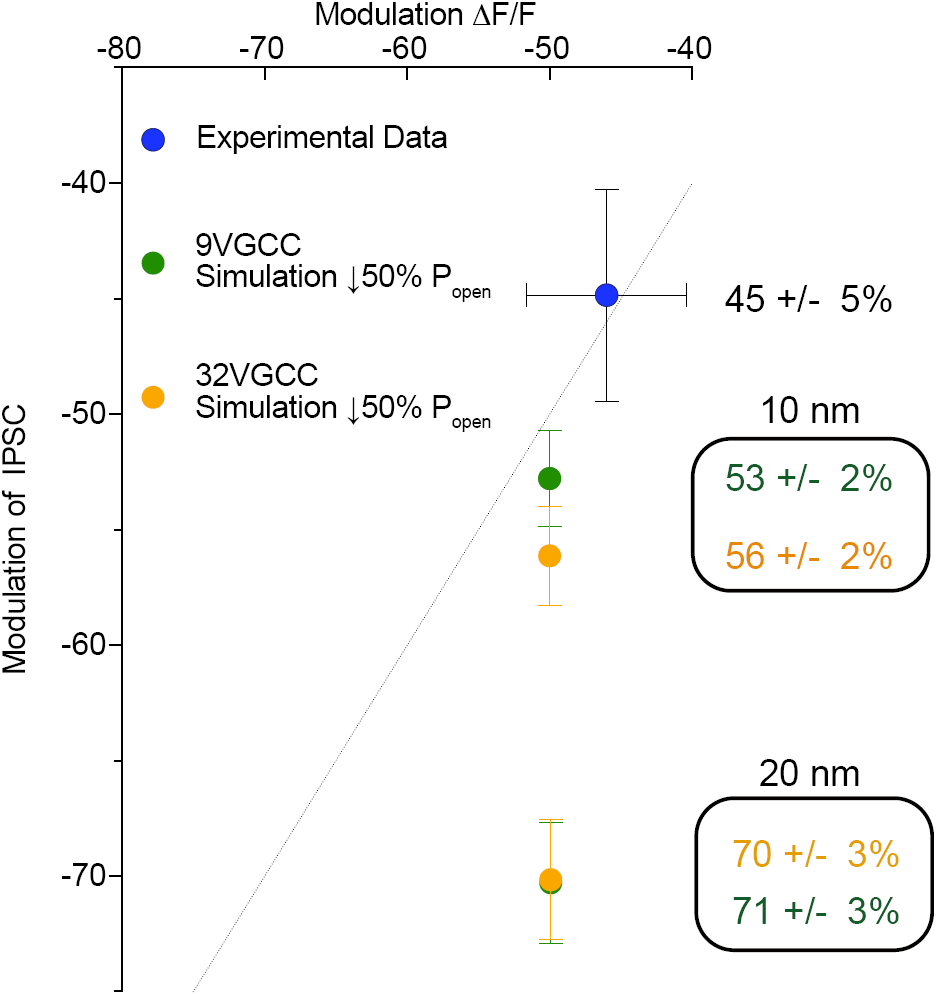
Calcium channel cooperativity for SC model. Impact of 50% decrease open probability of VGCCs in estimated P_v_ for PRM (Figure 6) using a cluster size of either 9 or 32 VGCCs. The release sensor was located at 10 or 20 nm from the edge of the cluster of VGCCs. Dashed line indicates linear relationship between modulation calcium entry and variation in P_v_. Model simulation predicted that at a coupling distance of 10 nm, decrease in calcium should be close to linear closely matching experimentally obtained data using baclofen (blue).

**Table S1.**
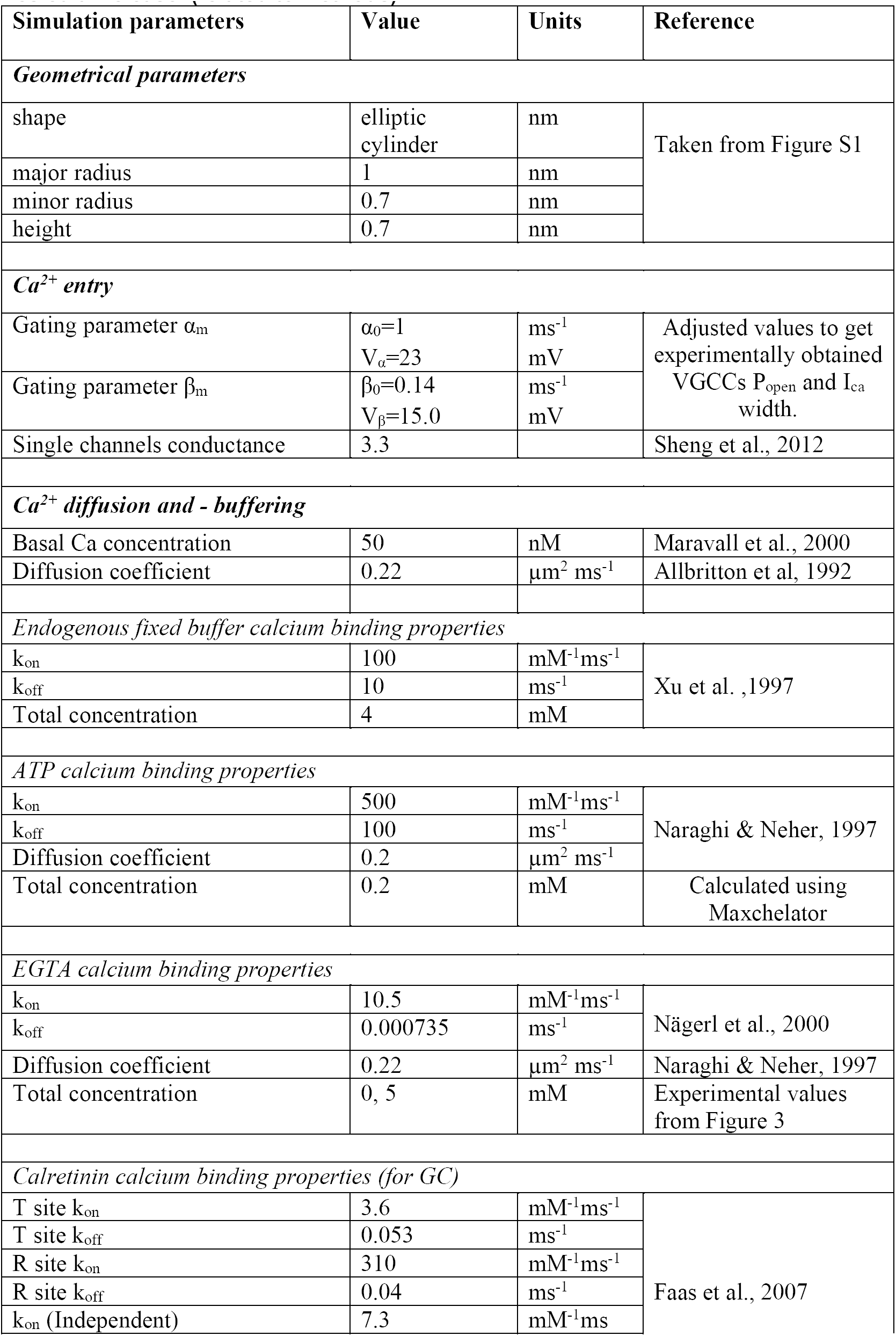
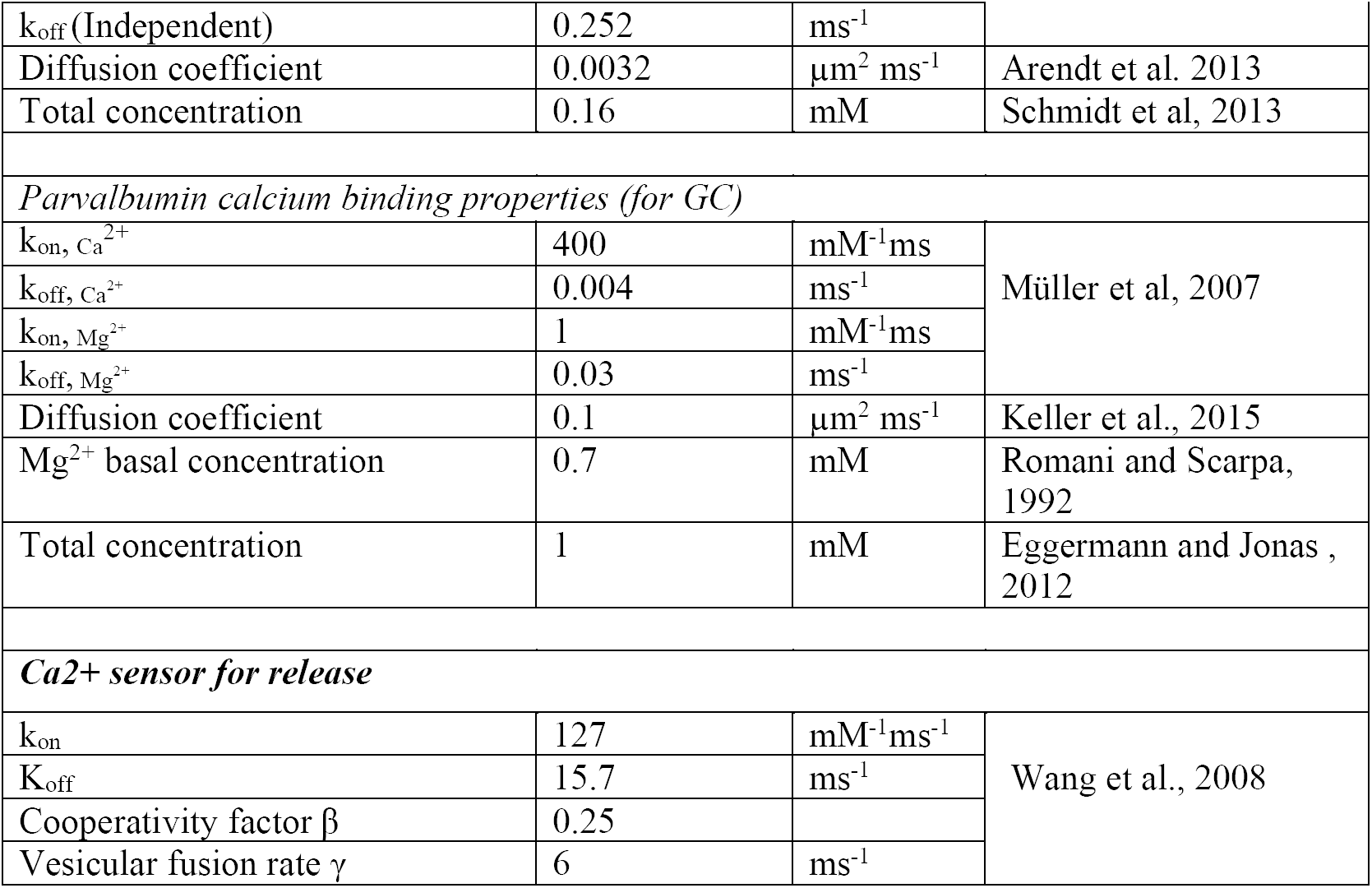
Model parameters for stochastic simulations of Calcium dynamics and Vesicular release. (related to Methods)

**Table S2.** Parameters of cluster generation for Figure S4 (related to Methods)

| $N_c$ | $N_p$ | $d_{cl}$ | $d_{pmin}$ | $d_{pmax}$ |
| --- | --- | --- | --- | --- |
| 5 | 36 | 130 | 5.3 | 95 |
| 20 | 9 | 60 | 5.3 | 45 |
| 60 | 3 | 25 | 5.3 | 20 |

## References

Abbott, L.F., and Regehr, W.G. (2004). Synaptic computation. Nature 431, 796–803.

Abrahamsson, T., Cathala, L., Matsui, K., Shigemoto, R., and Digregorio, D.A. (2012). Thin dendrites of cerebellar interneurons confer sublinear synaptic integration and a gradient of short-term plasticity. Neuron 73, 1159–1172.

Arai, I., and Jonas, P. (2014). Nanodomain coupling explains Ca(2)(+) independence of transmitter release time course at a fast central synapse. Elife 3.

Atwood, H.L., and Karunanithi, S. (2002). Diversification of synaptic strength: presynaptic elements. Nat Rev Neurosci 3, 497–516.

Augustin, I., Korte, S., Rickmann, M., Kretzschmar, H.A., Sudhof, T.C., Herms, J.W., and Brose, N. (2001). The cerebellum-specific Munc13 isoform Munc13-3 regulates cerebellar synaptic transmission and motor learning in mice. J Neurosci 21, 10–17.

Augustine, G.J. (1990). Regulation of transmitter release at the squid giant synapse by presynaptic delayed rectifier potassium current. J Physiol 431, 343–364.

Baddeley, A., Diggle, P.J., Hardegen, A., Lawrence, T., Milne, R.K. and Nair, G. (2014). On tests of spatial pattern based on simulation envelopes. Ecological Monographs 84, 477–489.

Baur, D., Bornschein, G., Althof, D., Watanabe, M., Kulik, A., Eilers, J., and Schmidt, H. (2015). Developmental tightening of cerebellar cortical synaptic influx-release coupling. J Neurosci 35, 1858–1871.

Besag, J. (1977). Contribution to the discussion of Dr Ripley’s paper. Journal of the Royal Statistical Society Series B 39, 193-195.

Biederer, T., Kaeser, P.S., and Blanpied, T.A. (2017). Transcellular Nanoalignment of Synaptic Function. Neuron 96, 680–696.

Bohme, M.A., Beis, C., Reddy-Alla, S., Reynolds, E., Mampell, M.M., Grasskamp, A.T., Lutzkendorf, J., Bergeron, D.D., Driller, J.H., Babikir, H., et al. (2016). Active zone scaffolds differentially accumulate Unc13 isoforms to tune Ca2+ channel-vesicle coupling. Nat Neurosci.

Borst, J.G., and Sakmann, B. (1998). Calcium current during a single action potential in a large presynaptic terminal of the rat brainstem. Journal of Physiology 506, 143–157.

Breustedt, J., Gundlfinger, A., Varoqueaux, F., Reim, K., Brose, N., and Schmitz, D. (2010). Munc13-2 differentially affects hippocampal synaptic transmission and plasticity. Cereb Cortex 20, 1109–1120.

Bucurenciu, I., Bischofberger, J., and Jonas, P. (2010). A small number of open Ca2+ channels trigger transmitter release at a central GABAergic synapse. Nat Neurosci 13, 19–21.

Bucurenciu, I., Kulik, A., Schwaller, B., Frotscher, M., and Jonas, P. (2008). Nanodomain coupling between Ca2+ channels and Ca2+ sensors promotes fast and efficient transmitter release at a cortical GABAergic synapse. Neuron 57, 536–545.

Calloway, N., Gouzer, G., Xue, M., and Ryan, T.A. (2015). The active-zone protein Munc13 controls the use-dependence of presynaptic voltage-gated calcium channels. Elife 4.

Chabrol, F.P., Arenz, A., Wiechert, M.T., Margrie, T.W., and DiGregorio, D.A. (2015). Synaptic diversity enables temporal coding of coincident multisensory inputs in single neurons. Nature neuroscience 18, 718–727.

Chadderton, P., Schaefer, A.T., Williams, S.R., and Margrie, T.W. (2014). Sensory-evoked synaptic integration in cerebellar and cerebral cortical neurons. Nature reviews Neuroscience 15, 71–83.

Chen, Z., Das, B., Nakamura, Y., DiGregorio, D.A., and Young, S.M., Jr. (2015). Ca2+ channel to synaptic vesicle distance accounts for the readily releasable pool kinetics at a functionally mature auditory synapse. The Journal of neuroscience: the official journal of the Society for Neuroscience 35, 2083–2100.

Datyner, N.B., and Gage, P.W. (1980). Phasic secretion of acetylcholine at a mammalian neuromuscular junction. J Physiol (Lond) 303, 299–314.

Delvendahl, I., Jablonski, L., Baade, C., Matveev, V., Neher, E., and Hallermann, S. (2015). Reduced endogenous Ca2+ buffering speeds active zone Ca2+ signaling. Proc Natl Acad Sci U S A 112, E3075–3084.

Deng, L., Kaeser, P.S., Xu, W., and Sudhof, T.C. (2011). RIM proteins activate vesicle priming by reversing autoinhibitory homodimerization of Munc13. Neuron 69, 317–331.

Diaz-Quesada, M., Martini, F.J., Ferrati, G., Bureau, I., and Maravall, M. (2014). Diverse thalamocortical short-term plasticity elicited by ongoing stimulation. J Neurosci 34, 515–526.

Dittman, J.S., Kreitzer, A.C., and Regehr, W.G. (2000). Interplay between facilitation, depression, and residual calcium at three presynaptic terminals. J Neurosci 20, 1374–1385.

Dittman, J.S., and Regehr, W.G. (1996). Contributions of calcium-dependent and calcium-independent mechanisms to presynaptic inhibition at a cerebellar synapse. J Neurosci 16, 1623–1633.

Dittrich, M., Pattillo, J.M., King, J.D., Cho, S., Stiles, J.R., and Meriney, S.D. (2013). An excess-calcium-binding-site model predicts neurotransmitter release at the neuromuscular junction. Biophys J 104, 2751–2763.

Eggermann, E., Bucurenciu, I., Goswami, S.P., and Jonas, P. (2012). Nanodomain coupling between Ca(2) channels and sensors of exocytosis at fast mammalian synapses. Nat Rev Neurosci 13, 7–21.

Eggermann, E., and Jonas, P. (2011). How the ‘slow’ Ca(2+) buffer parvalbumin affects transmitter release in nanodomain-coupling regimes. Nat Neurosci 15, 20–22.

Eltes, T., Kirizs, T., Nusser, Z., and Holderith, N. (2017). Target Cell Type-Dependent Differences in Ca2+ Channel Function Underlie Distinct Release Probabilities at Hippocampal Glutamatergic Terminals. J Neurosci 37, 1910–1924.

Ermolyuk, Y.S., Alder, F.G., Surges, R., Pavlov, I.Y., Timofeeva, Y., Kullmann, D.M., and Volynski, K.E. (2013). Differential triggering of spontaneous glutamate release by P/Q-, N-and R-type Ca(2+) channels. Nat Neurosci 16, 1754–1763.

Ester, M., Kriegel, H.-P., Sander, J., and Xu, X. (1996). A density-based algorithm for discovering clusters in large spatial databases with noise. Paper presented at: Kdd.

Fedchyshyn, M.J., and Wang, L.Y. (2005). Developmental transformation of the release modality at the calyx of held synapse. J Neurosci 25, 4131–4140.

Glebov, O.O., Jackson, R.E., Winterflood, C.M., Owen, D.M., Barker, E.A., Doherty, P., Ewers, H., and Burrone, J. (2017). Nanoscale Structural Plasticity of the Active Zone Matrix Modulates Presynaptic Function. Cell Rep 18, 2715–2728.

Grande, G., and Wang, L.Y. (2011). Morphological and functional continuum underlying heterogeneity in the spiking fidelity at the calyx of Held synapse in vitro. J Neurosci 31, 13386–13399.

Grauel, M.K., Maglione, M., Reddy-Alla, S., Willmes, C.G., Brockmann, M.M., Trimbuch, T., Rosenmund, T., Pangalos, M., Vardar, G., Stumpf, A., et al. (2016). RIM-binding protein 2 regulates release probability by fine-tuning calcium channel localization at murine hippocampal synapses. Proc Natl Acad Sci U S A 113, 11615–11620.

Han, Y., Kaeser, P.S., Sudhof, T.C., and Schneggenburger, R. (2011). RIM determines Ca(2)+ channel density and vesicle docking at the presynaptic active zone. Neuron 69, 304–316.

Hanisch, K.H.S.D. (1979). Formulas for second-order analysis of marked point processes, Mathematische Operationsforschung und Statistik, Series Statistics 10, 555–560.

Harlow, M.L., Ress, D., Stoschek, A., Marshall, R.M., and McMahan, U.J. (2001). The architecture of active zone material at the frog’s neuromuscular junction. Nature 409, 479–484.

Holderith, N., Lorincz, A., Katona, G., Rozsa, B., Kulik, A., Watanabe, M., and Nusser, Z. (2012). Release probability of hippocampal glutamatergic terminals scales with the size of the active zone. Nat Neurosci 15, 988–997.

Hruska, M., Henderson, N., Le Marchand, S.J., Jafri, H., and Dalva, M.B. (2018). Synaptic nanomodules underlie the organization and plasticity of spine synapses. Nat Neurosci.

Isope, P., and Barbour, B. (2002). Properties of unitary granule cell-->Purkinje cell synapses in adult rat cerebellar slices. J Neurosci 22, 9668–9678.

Jackman, S.L., and Regehr, W.G. (2017). The Mechanisms and Functions of Synaptic Facilitation. Neuron 94, 447–464.

Kaeser, P.S., Deng, L., Wang, Y., Dulubova, I., Liu, X., Rizo, J., and Sudhof, T.C. (2011). RIM proteins tether Ca2+ channels to presynaptic active zones via a direct PDZ-domain interaction. Cell 144, 282–295.

Katz, B. (1969). The Release of Neural Transmitter Substances (Liverpool: Liverpool University Press).

Keller, D., Babai, N., Kochubey, O., Han, Y., Markram, H., Schurmann, F., and Schneggenburger, R. (2015). An Exclusion Zone for Ca2+ Channels around Docked Vesicles Explains Release Control by Multiple Channels at a CNS Synapse. PLoS Comput Biol 11, e1004253.

Kirizs, T., Kerti-Szigeti, K., Lorincz, A., and Nusser, Z. (2014). Distinct axo-somato-dendritic distributions of three potassium channels in CA1 hippocampal pyramidal cells. Eur J Neurosci 39, 1771–1783.

Kiskowski, M.A., Hancock, J.F., and Kenworthy, A.K. (2009). On the use of Ripley’s K-function and its derivatives to analyze domain size. Biophys J 97, 1095–1103.

Koester, H.J., and Johnston, D. (2005). Target cell-dependent normalization of transmitter release at neocortical synapses. Science 308, 863–866.

Korber, C., and Kuner, T. (2016). Molecular Machines Regulating the Release Probability of Synaptic Vesicles at the Active Zone. Front Synaptic Neurosci 8, 5.

Kusch, V., Bornschein, G., Loreth, D., Bank, J., Jordan, J., Baur, D., Watanabe, M., Kulik, A., Heckmann, M., Eilers, J., et al. (2018). Munc13-3 Is Required for the Developmental Localization of Ca(2+) Channels to Active Zones and the Nanopositioning of Cav2.1 Near Release Sensors. Cell Rep 22, 1965–1973.

Laghaei, R., Ma, J., Tarr, T.B., Homan, A.E., Kelly, L., Tilvawala, M.S., Vuocolo, B.S., Rajasekaran, H.P., Meriney, S.D., and Dittrich, M. (2018). Transmitter release site organization can predict synaptic function at the neuromuscular junction. J Neurophysiol 119, 1340–1355.

Lenkey, N., Kirizs, T., Holderith, N., Mate, Z., Szabo, G., Vizi, E.S., Hajos, N., and Nusser, Z. (2015). Tonic endocannabinoid-mediated modulation of GABA release is independent of the CB1 content of axon terminals. Nature communications 6, 6557.

Liu, K.S., Siebert, M., Mertel, S., Knoche, E., Wegener, S., Wichmann, C., Matkovic, T., Muhammad, K., Depner, H., Mettke, C., et al. (2011). RIM-binding protein, a central part of the active zone, is essential for neurotransmitter release. Science 334, 1565–1569.

Luo, F., Dittrich, M., Cho, S., Stiles, J.R., and Meriney, S.D. (2015). Transmitter release is evoked with low probability predominately by calcium flux through single channel openings at the frog neuromuscular junction. J Neurophysiol 113, 2480–2489.

Matveev, V., Bertram, R., and Sherman, A. (2011). Calcium cooperativity of exocytosis as a measure of Ca(2)+ channel domain overlap. Brain Res 1398, 126–138.

Miki, T., Kaufmann, W.A., Malagon, G., Gomez, L., Tabuchi, K., Watanabe, M., Shigemoto, R., and Marty, A. (2017). Numbers of presynaptic Ca(2+) channel clusters match those of functionally defined vesicular docking sites in single central synapses. Proc Natl Acad Sci U S A 114, E5246–E5255.

Millar, A.G., Bradacs, H., Charlton, M.P., and Atwood, H.L. (2002). Inverse relationship between release probability and readily releasable vesicles in depressing and facilitating synapses. J Neurosci 22, 9661–9667.

Mintz, I.M., Sabatini, B.L., and Regehr, W.G. (1995). Calcium control of transmitter release at a cerebellar synapse. Neuron 15, 675–688.

Mittmann, W., Koch, U., and Hausser, M. (2005). Feed-forward inhibition shapes the spike output of cerebellar Purkinje cells. J Physiol 563, 369–378.

Nagwaney, S., Harlow, M.L., Jung, J.H., Szule, J.A., Ress, D., Xu, J., Marshall, R.M., and McMahan, U.J. (2009). Macromolecular connections of active zone material to docked synaptic vesicles and presynaptic membrane at neuromuscular junctions of mouse. J Comp Neurol 513, 457–468.

Nakamura, Y., Harada, H., Kamasawa, N., Matsui, K., Rothman, J.S., Shigemoto, R., Silver, R.A., DiGregorio, D.A., and Takahashi, T. (2015). Nanoscale distribution of presynaptic Ca(2+) channels and its impact on vesicular release during development. Neuron 85, 145–158.

Nakamura, Y., Reva, M., and DiGregorio, D.A. (2018). Variations in Ca(2+) Influx Can Alter Chelator-Based Estimates of Ca(2+) Channel-Synaptic Vesicle Coupling Distance. J Neurosci 38, 3971–3987.

Neef, J., Urban, N.T., Ohn, T.L., Frank, T., Jean, P., Hell, S.W., Willig, K.I., and Moser, T. (2018). Quantitative optical nanophysiology of Ca(2+) signaling at inner hair cell active zones. Nat Commun 9, 290.

Nielsen, T.A., DiGregorio, D.A., and Silver, R.A. (2004). Modulation of glutamate mobility reveals the mechanism underlying slow-rising AMPAR EPSCs and the diffusion coefficient in the synaptic cleft. Neuron 42, 757–771.

Nusser, Z., Cull-Candy, S., and Farrant, M. (1997). Differences in synaptic GABA(A) receptor number underlie variation in GABA mini amplitude. Neuron 19, 697–709.

Nusser, Z., Sieghart, W., Benke, D., Fritschy, J.M., and Somogyi, P. (1996). Differential synaptic localization of two major gamma-aminobutyric acid type A receptor alpha subunits on hippocampal pyramidal cells. Proc Natl Acad Sci U S A 93, 11939–11944.

Pangrsic, T., Gabrielaitis, M., Michanski, S., Schwaller, B., Wolf, F., Strenzke, N., and Moser, T. (2015). EF-hand protein Ca2+ buffers regulate Ca2+ influx and exocytosis in sensory hair cells. Proc Natl Acad Sci U S A 112, E1028–1037.

Pulido, C., Trigo, F.F., Llano, I., and Marty, A. (2015). Vesicular release statistics and unitary postsynaptic current at single GABAergic synapses. Neuron 85, 159–172.

Rangaraju, V., Calloway, N., and Ryan, T.A. (2014). Activity-driven local ATP synthesis is required for synaptic function. Cell 156, 825–835.

Reddy-Alla, S., Bohme, M.A., Reynolds, E., Beis, C., Grasskamp, A.T., Mampell, M.M., Maglione, M., Jusyte, M., Rey, U., Babikir, H., et al. (2017). Stable Positioning of Unc13 Restricts Synaptic Vesicle Fusion to Defined Release Sites to Promote Synchronous Neurotransmission. Neuron 95, 1350–1364 e1312.

Ripley, B.D. (1977). Modelling spatial patterns. Journal of the Royal Statistical Society Series B 39, 172–212.

Robitaille, R., Adler, E.M., and Charlton, M.P. (1990). Strategic location of calcium channels at transmitter release sites of frog neuromuscular synapses. Neuron 5, 773–779.

Rothman, J.S., and Silver, R.A. (2018). NeuroMatic: An Integrated Open-Source Software Toolkit for Acquisition, Analysis and Simulation of Electrophysiological Data. Front Neuroinform 12, 14.

Rowan, M.J., DelCanto, G., Yu, J.J., Kamasawa, N., and Christie, J.M. (2016). Synapse-Level Determination of Action Potential Duration by K(+) Channel Clustering in Axons. Neuron 91, 370–383.

Rozov, A., Burnashev, N., Sakmann, B., and Neher, E. (2001). Transmitter release modulation by intracellular Ca2+ buffers in facilitating and depressing nerve terminals of pyramidal cells in layer 2/3 of the rat neocortex indicates a target cell-specific difference in presynaptic calcium dynamics. J Physiol 531, 807–826.

Sabatini, B.L., and Regehr, W.G. (1997). Control of neurotransmitter release by presynaptic waveform at the granule cell to Purkinje cell synapse. Journal of Neuroscience 17, 3425–3435.

Sabatini, B.L., and Regehr, W.G. (1998). Optical measurement of presynaptic calcium currents. Biophys J 74, 1549–1563.

Sabatini, B.L., and Svoboda, K. (2000). Analysis of calcium channels in single spines using optical fluctuation analysis. Nature 408, 589–593.

Sakamoto, H., Ariyoshi, T., Kimpara, N., Sugao, K., Taiko, I., Takikawa, K., Asanuma, D., Namiki, S., and Hirose, K. (2018). Synaptic weight set by Munc13-1 supramolecular assemblies. Nat Neurosci 21, 41–49.

Sargent, P.B., Saviane, C., Nielsen, T.A., DiGregorio, D.A., and Silver, R.A. (2005). Rapid vesicular release, quantal variability, and spillover contribute to the precision and reliability of transmission at a glomerular synapse. J Neurosci 25, 8173–8187.

Schmidt, H., Brachtendorf, S., Arendt, O., Hallermann, S., Ishiyama, S., Bornschein, G., Gall, D., Schiffmann, S.N., Heckmann, M., and Eilers, J. (2013). Nanodomain coupling at an excitatory cortical synapse. Curr Biol 23, 244–249.

Scimemi, A., and Diamond, J.S. (2012). The number and organization of Ca2+ channels in the active zone shapes neurotransmitter release from Schaffer collateral synapses. J Neurosci 32, 18157–18176.

Sudhof, T.C. (2012). The presynaptic active zone. Neuron 75, 11–25.

Szoboszlay, M., Kirizs, T., and Nusser, Z. (2017). Objective quantification of nanoscale protein distributions. Sci Rep 7, 15240.

Tang, A.H., Chen, H., Li, T.P., Metzbower, S.R., MacGillavry, H.D., and Blanpied, T.A. (2016). A trans-synaptic nanocolumn aligns neurotransmitter release to receptors. Nature 536, 210–214.

Tian, T., Harding, A., Inder, K., Plowman, S., Parton, R.G., and Hancock, J.F. (2007). Plasma membrane nanoswitches generate high-fidelity Ras signal transduction. Nat Cell Biol 9, 905–914.

Turecek, J., and Regehr, W.G. (2018). Synaptotagmin 7 Mediates Both Facilitation and Asynchronous Release at Granule Cell Synapses. J Neurosci 38, 3240–3251.

Valera, A.M., Doussau, F., Poulain, B., Barbour, B., and Isope, P. (2012). Adaptation of granule cell to Purkinje cell synapses to high-frequency transmission. J Neurosci 32, 3267–3280.

Van der Kloot, W. (1988). Estimating the timing of quantal releases during end-plate currents at the frog neuromuscular junction. Journal of Physiology 402, 595–603.

Vyleta, N.P., and Jonas, P. (2014). Loose coupling between Ca2+ channels and release sensors at a plastic hippocampal synapse. Science 343, 665–670.

Wang, L.Y., Neher, E., and Taschenberger, H. (2008). Synaptic vesicles in mature calyx of Held synapses sense higher nanodomain calcium concentrations during action potential-evoked glutamate release. J Neurosci 28, 14450–14458.

Wong, A.B., Rutherford, M.A., Gabrielaitis, M., Pangrsic, T., Gottfert, F., Frank, T., Michanski, S., Hell, S., Wolf, F., Wichmann, C., et al. (2014). Developmental refinement of hair cell synapses tightens the coupling of Ca2+ influx to exocytosis. EMBO J 33, 247–264.

Xu-Friedman, M.A., Harris, K.M., and Regehr, W.G. (2001). Three-dimensional comparison of ultrastructural characteristics at depressing and facilitating synapses onto cerebellar Purkinje cells. J Neurosci 21, 6666–6672.

